# Neural mechanisms of parasite-induced summiting behavior in “zombie” *Drosophila*

**DOI:** 10.1101/2022.12.01.518723

**Authors:** Carolyn Elya, Danylo Lavrentovich, Emily Lee, Cassandra Pasadyn, Jasper Duval, Maya Basak, Valerie Saykina, Benjamin de Bivort

## Abstract

For at least two centuries, scientists have been enthralled by the “zombie” behaviors induced by mind-controlling parasites. Despite this interest, the mechanistic bases of these uncanny processes have remained mostly a mystery. Here, we leverage the recently established *Entomophthora muscae*-*Drosophila melanogaster* “zombie fly” system to reveal the molecular and cellular underpinnings of summit disease, a manipulated behavior evoked by many fungal parasites. Using a new, high-throughput behavior assay to measure summiting, we discovered that summiting behavior is characterized by a burst of locomotion and requires the host circadian and neurosecretory systems, specifically DN1p circadian neurons, pars intercerebralis to corpora allata projecting (PI-CA) neurons and corpora allata (CA), who are solely responsible for juvenile hormone (JH) synthesis and release. Summiting is a fleeting phenomenon, posing a challenge for physiological and biochemical experiments requiring tissue from summiting flies. We addressed this with a machine learning classifier to identify summiting animals in real time. PI-CA neurons and CA appear to be intact in summiting animals, despite *E. muscae* cells invading the host brain, particularly in the superior medial protocerebrum (SMP), the neuropil that contains DN1p axons and PI-CA dendrites. The blood-brain barrier of flies late in their infection was significantly permeabilized, suggesting that factors in the hemolymph may have greater access to the central nervous system during summiting. Metabolomic analysis of hemolymph from summiting flies revealed differential abundance of several compounds compared to non-summiting flies. Transfusing the hemolymph of summiting flies into non-summiting recipients induced a burst of locomotion, demonstrating that factor(s) in the hemolymph likely cause summiting behavior. Altogether, our work reveals a neuro-mechanistic model for summiting wherein fungal cells perturb the fly’s hemolymph, activating the neurohormonal pathway linking clock neurons to juvenile hormone production in the CA, ultimately inducing locomotor activity in their host.

## Introduction

Many organisms infect animals and compel them to perform specific, often bizarre, behaviors that serve to promote their own fitness at the expense of their host. For example, “zombie ant” fungi of genus *Ophiocordyceps* compel their host carpenter ants to aberrantly leave the nest, wander away from established foraging trails, scale nearby stems or twigs, and, in their dying moments, clamp onto vegetation to ultimately perish in elevated positions (Hughes et al., 2011; Pontoppidan et al., 2009). Days later, a fungal stalk emerges from the dead ant’s pronotum, well poised to rain spores on the ants that forage below (Evans and Samson, 1984). But this is far from the only example: jewel wasps that subdue cockroaches (Gal and Libersat, 2010), protozoans that suppress a rodent’s fear of cat odors (Vyas et al., 2007), and worms that drive crickets to leap to watery deaths are all examples of parasites hijacking host behavior (Thomas et al., 2002).

One of the most frequently encountered behavior manipulations in parasitized insects is summit disease (also referred to as tree-top disease or Wipfelkrankheit) (Hofmann, 1891). Summit disease is induced by diverse parasites, ranging from viruses to fungi to trematodes, and affects a broad range of insect species, including ants, beetles, crickets, caterpillars, and flies (Goulson, 1997; Hughes et al., 2011; Krasnoff et al., 1995; Loos-Frank and Zimmermann, 1976; Pickford and Riegert, 1964; Steinkraus et al., 2017). The most consistently reported symptom of summit disease is elevation prior to death (Evans, 1989; Lovett et al., 2020; Roy et al., 2006). This positioning advantages the parasite by either making the spent host more conspicuous, and therefore likely to be consumed by the next host in its life cycle (e.g., *Dicrocoelium dendriticum*-infected ants; Martín-Vega et al., 2018), or by positioning the spent host for optimal dispersal of infectious propagules (e.g., *Mamestra brassicae* nuclear polyhedrosis virus; Goulson, 1997).

Some of the deepest mechanistic understanding of parasite-induced summiting comes from nucleopolyhedroviruses (NPVs). Disrupting the *ecdysteroid uridine 5’-diphosphate* (*egt*) gene in NPVs of the moths *Lymantria dispar* or *Spodoptera exigua* prevents summiting in infected larvae (Han et al., 2015; Hoover et al., 2011). This effect is thought to occur via *egt*’s inactivation of the hormone 20-hydroxyecdysone and the resulting disruption of molting (O’Reilly and Miller, 1989). However, *egt* has been found to be dispensable for driving summit disease in other NPV-insect systems (Kokusho and Katsuma, 2021), suggesting there are undiscovered viral mechanisms driving summiting in NPV-infected hosts. On the host side, evidence in NPV-infected *L. dispar* and *Helicoverpa armigera* point to changes in the host phototactic pathway underlying summiting behavior (Bhattarai et al., 2018; Liu et al., 2022). Outside of NPVs, work in *Ophiocordyceps* suggests that the parasitic fungus may use enterotoxins and small secreted proteins to mediate end-of-life “zombie” behaviors (Beckerson et al., 2022; de Bekker et al., 2015; Will et al., 2020), potentially targeting host phototaxis (Andriolli et al., 2019), circadian rhythm, chemosensation, and locomotion (de Bekker et al., 2015; Trinh et al., 2021; Will et al., 2020).

*Entomophthora muscae* is a behavior-manipulating fungal pathogen that infects dipterans and elicits summit disease prior to host death (Graham-Smith, 1916; MacLeod et al., 1976). *E. muscae* infection begins when a fungal conidium (informally: spore) ejected from a dead host lands on a fly’s cuticle. The spore penetrates the cuticle and enters the hemolymph where it begins to replicate, first using the fat body (a tissue analogous to the liver and used for storing excess nutrients) as a food source (Brobyn and Wilding, 1983). When nutrients are exhausted, *E. muscae* elicits a stereotyped trio of behaviors to position its dying host for the next round of spore dispersal. The fly 1) summits (Graham-Smith, 1916), 2) extends its proboscis, which glues the fly in place via sticky, exuded secretions (Brobyn and Wilding, 1983), and finally, 3) the fly’s wings lift up and away from its dorsal abdomen, clearing the way for future spore dispersal (Elya et al., 2018; Krasnoff et al., 1995). Fungal structures (conidiophores) then emerge through the cuticle and forcefully eject infectious spores into the surrounding environment via a ballistic water cannon mechanism (de Ruiter et al., 2019). *E. muscae* kills flies with reliable circadian timing: flies die around sunset and exhibit their final bout of locomotion between 0-5 hours prior to lights off (Elya et al., 2018; Krasnoff et al., 1995). Circadian regulation is a common feature of fungal-induced summit disease: *Ophiocordyceps*-infected ants die around solar noon (Hughes et al., 2011), *Entomophaga grylli*-infected grasshoppers within a four hour window prior to sunset (Roffey, 1968), and *Erynia neoaphidis*- and *Entomophthora planchoniana*-infected aphids die most frequently around 8.5 and 14 hours after sunrise, respectively (Milner et al., 1984)

*E. muscae*-infected “zombie flies” have been known to the scientific literature for the last 167 years (Cohn, 1855), yet the mechanistic basis of their behavior manipulation is still a mystery. It is challenging to culture *E. muscae* in the laboratory and typical host species, like houseflies, lack experimental access. A strain of *E. muscae* that infects fruit flies was recently isolated and used to establish a laboratory-based “zombie fly” system in the tool-replete model organism *Drosophila melanogaster* (Elya et al., 2018), permitting investigation of the specific host mechanisms underlying manipulated behaviors.

The rich experimental toolkit of *D. melanogaster* has been used to decipher the mechanistic underpinnings of host-symbiont interactions ranging from mutualism to parasitism. For example, a mutant screen identified the Toll pathway as essential for *Drosophila*’s antiviral immune response (Zambon et al., 2005). Genetic access to specific neuronal populations allowed the identification of class IV neurons as mediating the larval escape response to oviposition by *Leptopilina boulardi* wasps (Robertson et al., 2013). It was recently shown that the gut bacterium *Lactobacillus brevis* alters fly octopaminergic pathways to drive an increase in locomotion (Schretter et al., 2018). Fruit flies have also been leveraged to investigate mechanisms of medically important parasites naturally vectored by other dipterans, including the protozoans *Plasmodium, Leishmania* and *Trypanosoma* (dos-Santos et al., 2015; Peltan et al., 2012; Tonk et al., 2019).

Here, we describe our progress using the zombie fruit fly system to unravel the mechanistic basis of summiting behavior. We first show that the hallmark of summiting behavior is an increase in locomotion beginning ∼2.5 hours before death. By combining the powerful fruit fly genetic tool kit with a custom high-throughput behavioral assay, we demonstrate that the fly circadian and neurosecretory systems—specifically DN1p clock neurons, pars intercerebralis projection neurons that innervate the corpora allata (PI-CA neurons), and the juvenile hormone-producing corpora allata—are essential components mediating summiting. Using a real-time machine learning classifier to identify the moment flies begin to summit, we were able to characterize the anatomy and physiology of summiting flies with temporal precision. We found that *E. muscae* specifically invades the brain region harboring DN1p axons and PI-CA dendrites. The hemolymph of summiting flies contains specific metabolites that, when transfused into recipient flies, induces summiting-like locomotion. Taken together, these experiments reveal that *E. muscae* uses hemolymph-borne factors, targets a specific neural circuit, and hijacks endogenous neurohormonal control of locomotion.

## Results

### A novel assay to measure summiting behavior

We first set out to develop an assay that would allow us to characterize the behavioral mechanisms of summit disease (Fig 1A). Given the variability in the day and exact time when flies die, and the unknown duration of summiting, our assay needed to accommodate continuous monitoring of flies over many hours. The assay also needed to allow flies to express behavior with respect to the direction of gravity. We also wanted to make sure our chambers provided enough space for flies to lift their wings without interference (Fig 1B). Each behavioral arena was 65 mm long along the main gravitational axis, 5 mm wide and 3.2 mm deep, and housed a single fly (Fig 1C). The bottom of the chamber was plugged with food to sustain flies over long periods of observation (24-96 hours). Four rows of 32 arenas each were fabricated in laser-cut acrylic trays, allowing us to measure the behavior (position along the main gravitational axis, referred to as “relative y position”, and overall speed) of 128 flies simultaneously. Trays and the imaging boxes that housed them were angled at 30 degrees (M. Reiser, personal communication) to provide the gravitactic gradient (Fig 1C).

**Figure 1.**
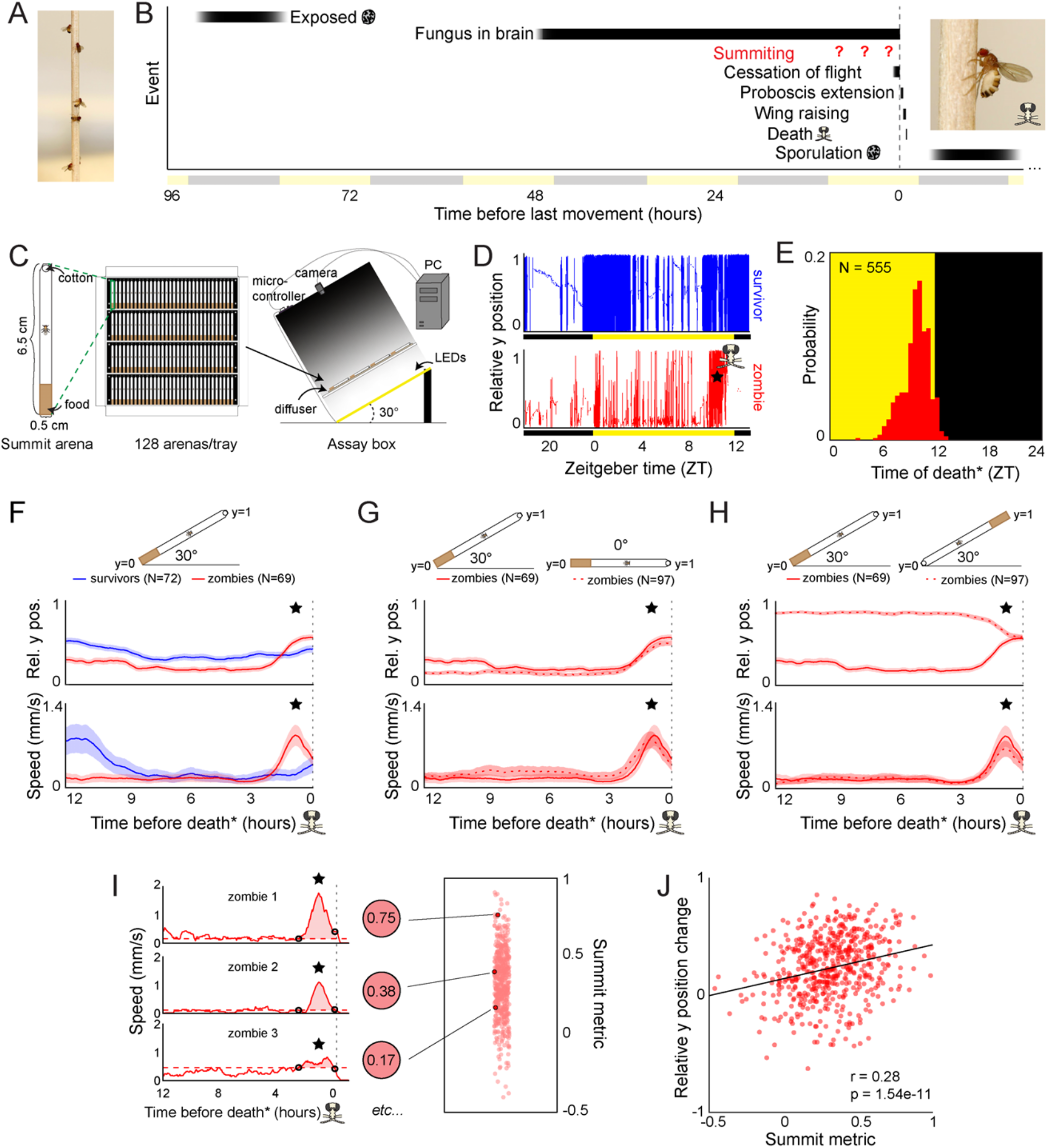
Behavioral signature of *E. muscae*-induced summiting in wild type flies. A) *E. muscae*-killed fruit flies that summited on a wooden dowel prior to death. B) Timeline of events relative to an *E. musace*-infected fly’s last movement (dashed line). See (Elya et al., 2018; Krasnoff et al., 1995). C) Summiting assay schematic. D) Example y position data for a typical survivor fly (top) and zombie (bottom). X-axis is Zeitgeber time (ZT), hours since lights were turned on. The fly “skull” indicates the manually-annotated time of zombie death (see Methods). Black and yellow bars indicate state of visible illumination. E) Distribution of time of death for Canton-S flies killed by *E. muscae*. Background color indicates state of visible illumination. F) Mean y position (middle) and mean speed (bottom) of survivor flies (blue) and zombie flies (red) housed in arenas angled at 30° with food at the bottom (schematic at top) during the 12 hours preceding the time of death. Here and in all other panels, shaded regions are +/- 1 standard error of the mean. Time of death for zombies was manually determined as the time of last movement from y position trace. Survivors did not die but were assigned fictive times of death from the distribution of zombie death times for comparability (see Methods). G) As in (F), but comparing zombies in standard arenas (30° with respect to gravity, same data as (F); solid lines) to zombies in flat arenas (0°; dashed lines). (H) As in (F) and (G), but comparing zombies in standard arenas (food at the bottom, same data as (F); solid lines) to zombies in arenas with food at the top (dashed lines). I) Speed versus time for three example Canton-S zombies (left) and their corresponding summit metrics (middle) outlined in black (right) amidst all Canton-S summit metric (right). Black circles denote the window of summiting behavior as determined from mean behavior of Canton-S zombie flies. Dashed red line indicates the mean speed in the hour preceding summiting (baseline speed). Summit metric is calculated as the integral of speed minus baseline in the summiting window (shaded region). J) Relative y position change versus summit metric for Canton-S zombies. Points are individual flies. Linear regression line in black; Pearson’s correlation r & p-value (upper left).

We first monitored *E. muscae*-exposed wild-type (Canton-S) flies. Experiments started no later than Zeitgeber time 20 (ZT20, i.e., 19 hours after the dark-to-light transition) on the day prior to their earliest possible death, until flies either succumbed to or survived their infection (ZT13 of day 4 to 7, depending on the experiment). After tracking, we manually assessed if each fly was alive or dead, and if the latter, whether it had sporulated. Henceforth, we will use the term “zombies” as a shorthand for *E. muscae*-exposed flies that perform fungus-induced behaviors before dying and sporulating. Sporulated flies were retroactively declared “zombies” and living flies “survivors.” Dead flies without signs of sporulation were excluded from further analysis. The time of zombie deaths was manually determined by the time of last movement (Fig 1D). As expected, wild-type flies killed by *E. muscae* tended to die in the evening (mean death time = ZT9:50 Fig 1E), but there was variability in the timing of death. 90% of all deaths occurred between ZT7 and ZT12.

### A burst of locomotion before death is a key signature of E. muscae-induced summiting

With our assay in its standard configuration (angled 30° with respect to gravity, food at the bottom), *E. muscae*-exposed survivors and zombies exhibited significantly different time-varying patterns in mean vertical position and mean speed in the final 12 hours before death (Fig 1F; survivors were randomly assigned a fictive time of death to enable this comparison). Survivor flies typically resided close to the center of the summit arena throughout tracking. In contrast, the average position of the zombie fly was near the bottom of the arena until approximately 2.5 hours before death when the average elevation increased, ultimately surpassing that of survivors. The difference between zombies and survivors in average speed over time was even more striking. Zombies maintained a low average speed (0.18 mm/s) until ∼2.5 hours before death when it increased substantially, peaking at 0.87 mm/s approximately one hour prior to death. In contrast, survivors exhibited high mean speed (∼0.8 mm/s) ∼12 h prior to the end of the experiment and a small increase in mean activity (0.22 m/s) ∼2h after the burst of zombie activity. These peaks of survivor activity correspond to the crepuscular peaks of activity expected in healthy flies.

Surprisingly, the average “elevation” and speed trajectories of zombie flies did not change in the absence of a gravitactic gradient (i.e., when the arena was laid flat, and the food was designated as the “bottom” of the arena) (Fig 1G). Flies resided near the food and exhibited low average speed (0.19 mm/s) until ∼2.5 hours prior to death, when speed peaked at 0.8 mm/s and flies had a mean position near the middle of the chamber. These patterns were largely statistically indistinguishable from those of the 30° experiment. When the chamber was angled at 30°, but with food at the top, average y position trends were essentially flipped, with flies on average residing near the top of the chamber until 2.5 hours prior to death, at which point they moved downward (Fig 1H). Notably, speed trends were statistically indistinguishable in this new configuration: flies still exhibited low average speed (0.15 mm/s) until ∼2.5 hours prior to death when they exhibited a marked increase in speed peaking at 0.66 mm/s ∼1 hour prior to death.

The burst of speed prior to death in zombie flies was specific to how they died. Unexposed flies that were killed by starvation (Fig 1-S1A) or desiccation (Fig 1-S1B) did not exhibit a burst of speed prior to death. In both cases, flies maintained a high average speed at 12 hours before death (2.2 mm/s and 2.9 mm/s, respectively) with the average speed of starved flies gradually declining over ∼5 hours before death. The mean speed of desiccated flies gradually increased from 12 to ∼3 hours before death, peaking at 4.85 mm/s, then exhibited a steady decline until death. Unlike zombie flies, starved or desiccated unexposed flies did not die at a specific time of day (Fig 1-S1C, S1D). These experiments suggest that an increase in speed ∼2.5 hours before death and dying at specific times are signatures of *E. muscae* mortality.

Average zombie y position appeared to be dictated by the location of food in our assay. Zombie flies began to reside closer to the food than survivors starting ∼24 hours prior to death in the food-at-the-top configuration (Fig 1-S1E). This behavior was dependent on the nutritive content of the food. When given a choice between sugar-containing and sugarless agar in a 0° assay, zombie flies tended to reside near the sugar-containing media before moving away ∼2.5 h prior to death (Fig 1-S1F). Providing food within the last 24 hours was necessary for the pre-death burst of locomotion: flies that were housed on sugarless media starting the day prior to death failed to exhibit a pre-death burst of locomotion (Fig 1-S1G) though still died with the expected circadian timing (Fig 1-S1H). These results suggest that flies are likely starving by late infection (Elya et al., 2018) and need access to sustenance to exhibit a final burst of locomotion during summiting.

A burst of locomotion will move flies, on average, away from the closed end of an arena, a consequence of that boundary condition. We were curious what would happen if flies were residing at food in the middle of an arena at the onset of summiting. We lengthened the arena and situated the food in the middle. As expected, in 0° arenas, zombie flies remained on average centered on the food prior to death (Fig 1-S1I). However, in 30° arenas, zombie flies moved on average slightly upward at the end of life (Fig 1-S1I). The distance that flies traveled during summiting did not differ between arenas angled at 0 to 30 degrees (Fig 1-S1J,K), indicating that the net upward motion of summiting in this condition could not be attributed to differences in activity.

Taken together, these experiments reveal a burst of speed in the final 2.5 hours before death as a key signature of *E. muscae*-induced summiting in our assay. We devised a simple metric, the summit metric (SM), to quantify the “summity-ness” of individual flies. SM is calculated as the integral of baseline-corrected speed over the summiting window. Three example speed traces for Canton-S flies and their corresponding SM values are shown in Fig 1I. As expected, there was a weak, positive correlation across individual flies between SM and change in y-position over summiting (Fig 1J). Comparing SM values across over 400 males and female Canton-S flies, we observed that, on average, males are moderately more “summity” (have 18% higher SM values) than females (Fig 1-S1L,M). However, this difference is dwarfed by interindividual variation in summiting, and since *E. muscae* infects both males and females in the wild, we opted to use mixed-sex experimental groups in subsequent experiments.

### Summiting behavior requires host circadian and neurosecretory pathways

With the understanding that a burst of activity shortly before death is the signature of summiting in this assay, we performed a screen to identify circuit and genetic components mediating summiting in the host fly. We adopted a candidate approach, focusing on neuromodulators and neuropeptides as well as neurons and genes that had been implicated in arousal and gravitaxis (Fig 2A-C). In neuron disruption experiments, we used tetanus toxin (TNT-E; a vesicle release blocker; Keller et al., 2002) to constitutively silence neurons. The specific pattern of TNT-E expression was determined by 103 different Gal4 drivers (Fig 2B, Table 1), with the effect size of each perturbation measured relative to a common heterozygous reagent (*UAS-TNT-E*/+) control. Similarly, we screened 101 lines targeting candidate genes, either by pan-neuronally reducing their expression via RNAi or testing mutant alleles (Figure 2C, Table 1). Effect sizes were estimated by comparing each line’s summiting metric to the heterozygous pan-neuronal driver (*R57C10-Gal4*/+) or wild-type control, respectively. In both the circuit and genetic screens we observed a range of effects on summiting from extreme impairment of the behavior (effect size -1) to rare amplification of summiting (effect size > 0). Most perturbations had effects that were not statistically distinguishable from zero.

**Table 1.**
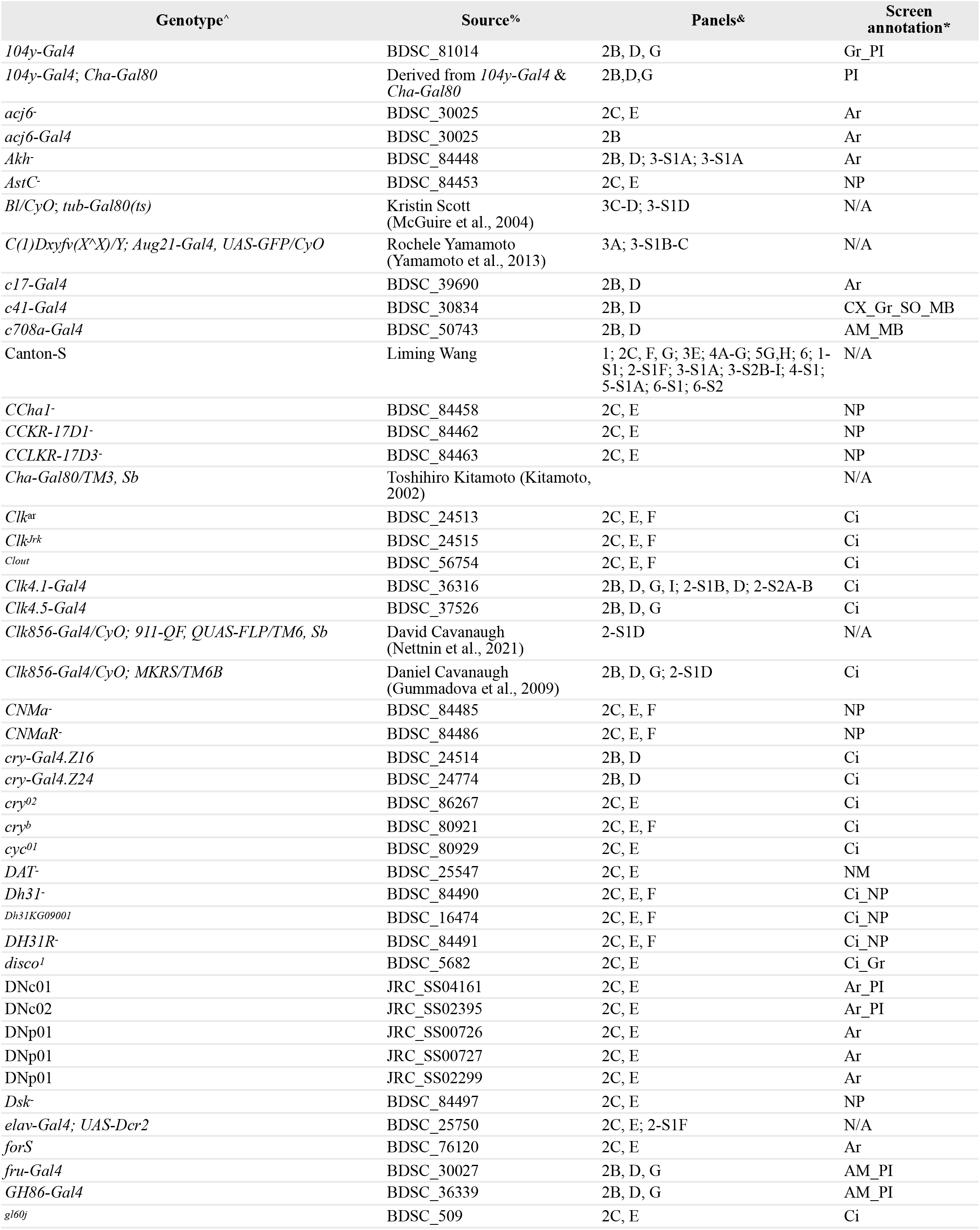
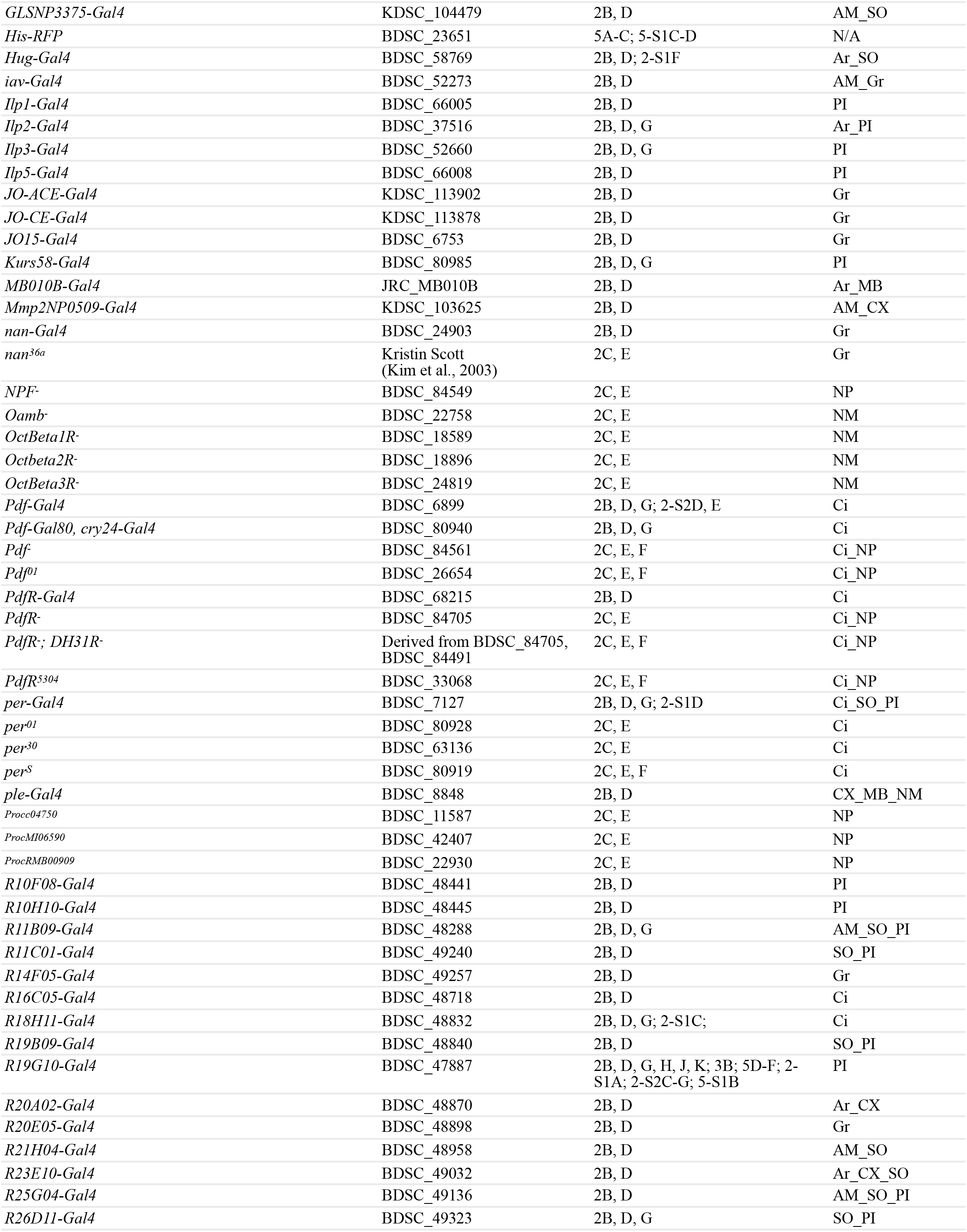
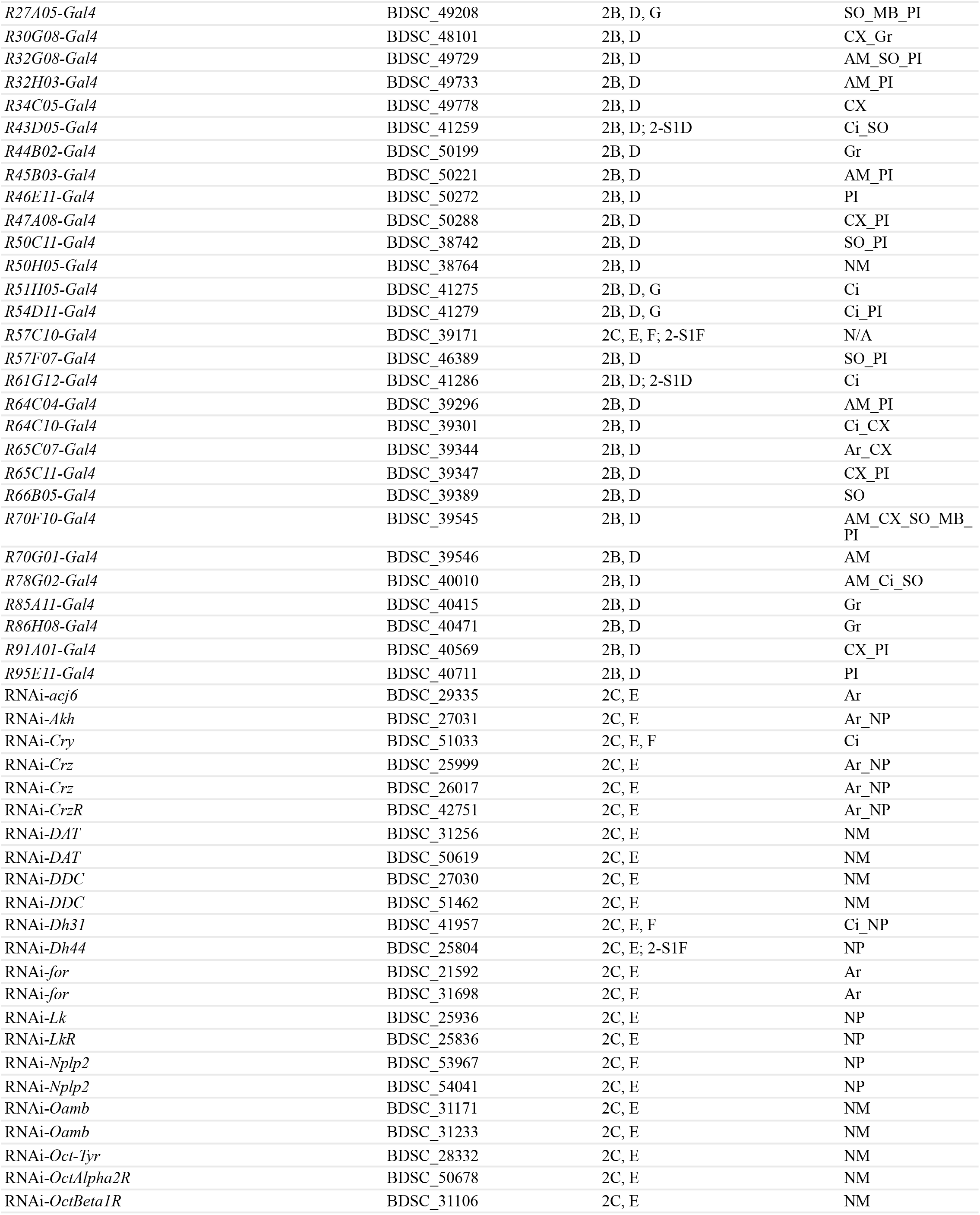
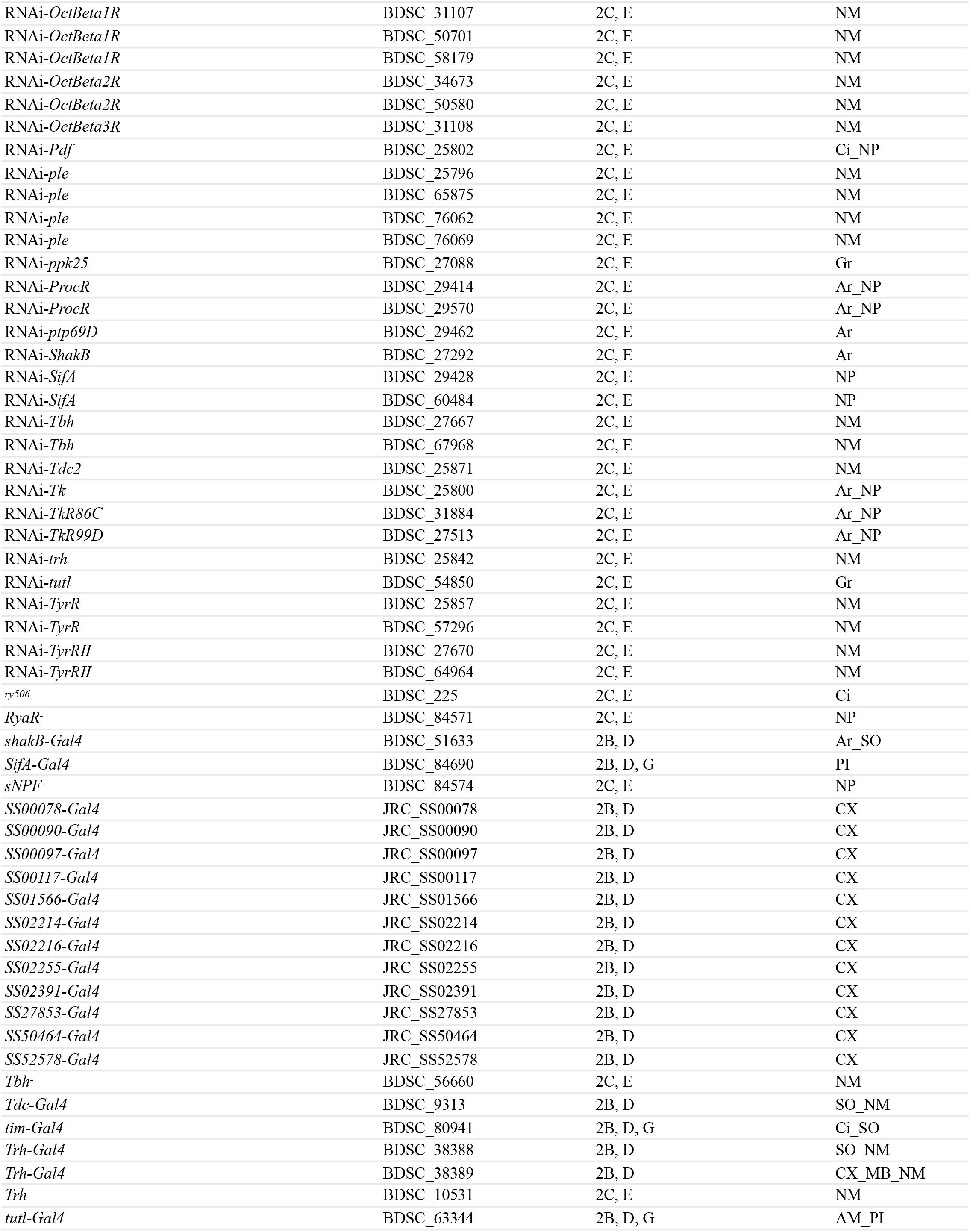
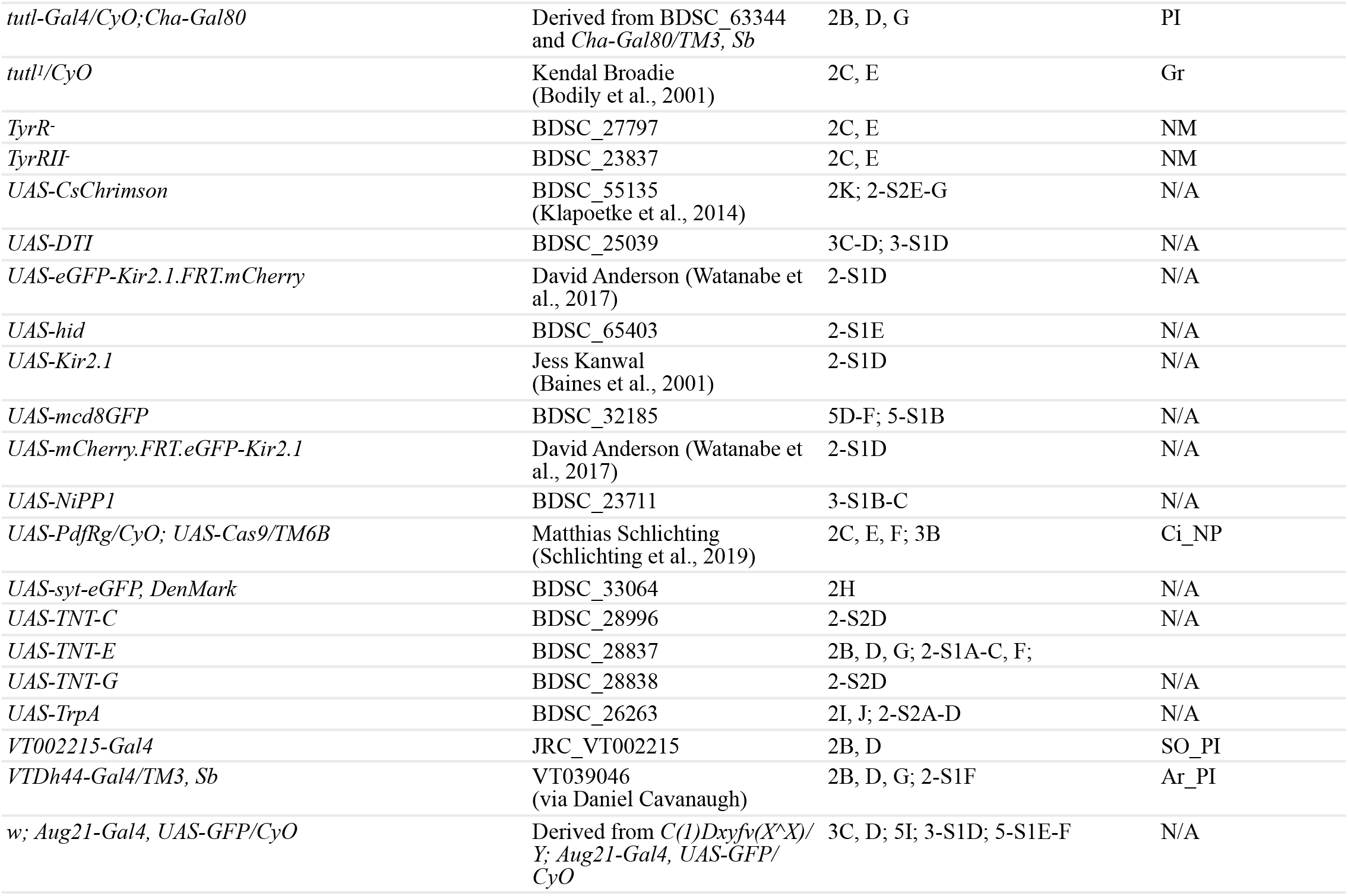
Fly strains used throughout this manuscript. ^ Genotypes are abbreviated for lines deposited at stock centers. % BDSC = Bloomington Drosophila Stock Center; KDSC = Kyoto Drosophila Stock Center; JRC = Janelia Research Campus & Lines are included in a panel if they were used for either experimental or control genotypes. * AM = AMMC, Ar = arousal, Ci = circadian, CX = central complex, Gr = gravitaxis, SO = subesophageal ganglion, MB = mushroom body, NM = neuromodulator & neurotransmitter, NP = neuropeptide, PI = pars intercerebralis.

**Figure 2.**
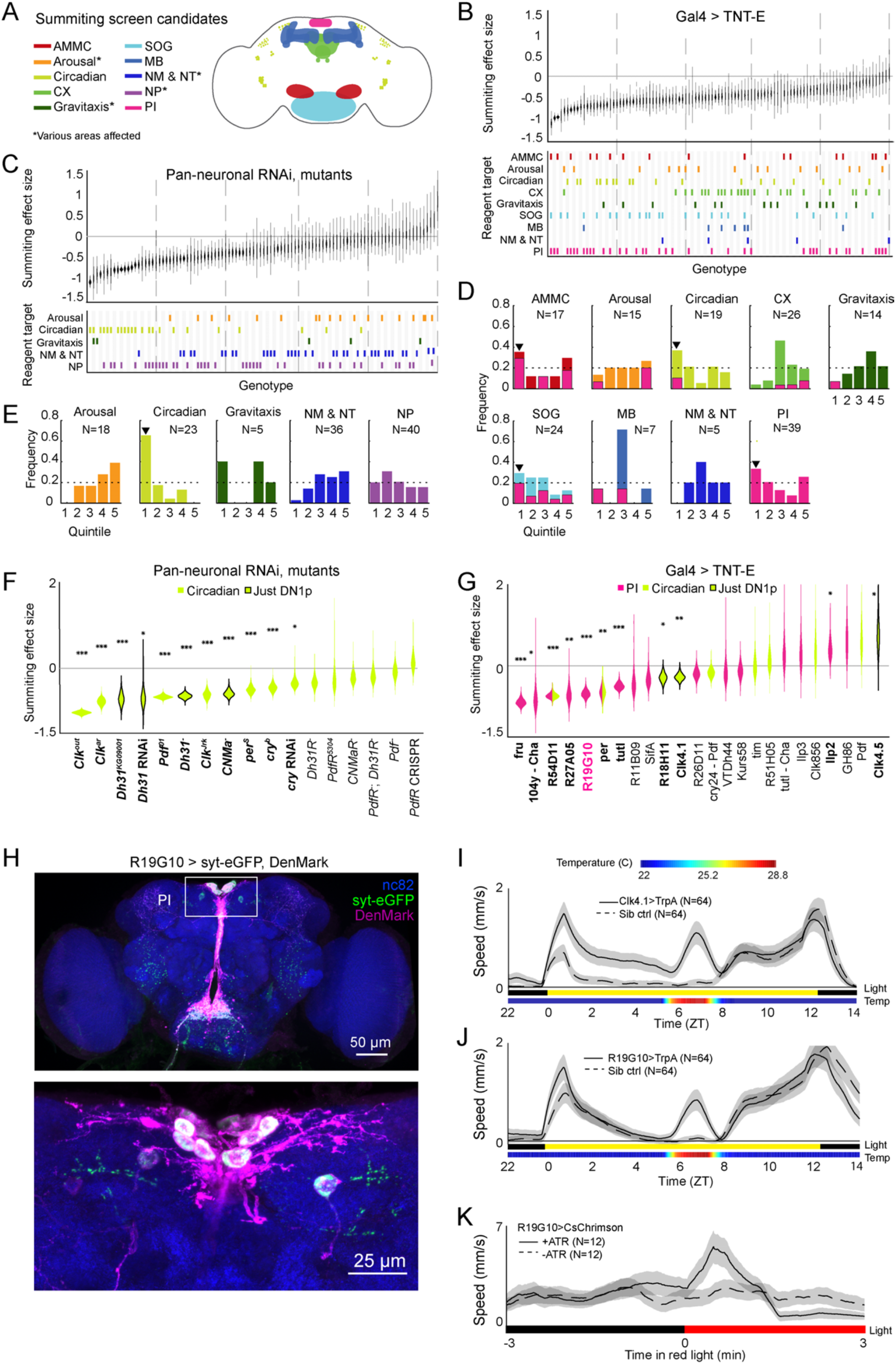
Identification of host circuit and genetic components involved in summiting behavior. A) Regions and pathways targeted in candidate screen. AMMC = antennal mechanosensory and motor center; CX = central complex; SOG = subesophageal ganglion; MB = mushroom body; NM & NT = neuromodulator or neurotransmitter; NP = neuropeptide; PI = pars intercerebralis. B & C) Effects of neuronal disruption (B) or gene knockdown or mutagenesis (C) on summiting. Above: Summiting effect size estimate distributions as estimated by bootstrapping. Experimental groups are ordered by mean effect (negative to positive). Below: gene function and brain region annotations associated with each screened reagent. See Table 1 for genotype and annotation details. Solid gray line indicates effect size of zero. Dashed vertical lines separate ranked data into quintiles. D & E) Frequency of annotations by quintile for B) and C), respectively. The number of lines screened (N) is indicated for each annotation. Dashed line indicates the frequency of annotation expected from a null, uniform distribution. Black arrowheads highlight annotations that are overrepresented in the first quintile. For D), pink overlays indicate portion of line annotations that are co-annotated for expression in the PI. F & G) Summiting effect size estimate distributions of disrupting specific circadian genes (F) or circadian and/or PI neurons (G) compared to genotype-matched controls. Lines are ordered by effect size. Pink indicates Gal4 expression in the PI, lime circadian Gal4 lines and genes, and black outlines expression only in DN1ps. Asterisks indicate statistically significant effects on summiting behavior by two-tailed t-test (* = p<0.05; ** = p<0.01; *** p<0.001). R19G10 is highlighted in pink to emphasize its subsequent use as the main PI reagent. See Table 2 for genotypes and matched controls. H) Maximum z-projections of brains showing pre- (synaptotagmin; syt-eGFP) and post- (DenMark) synaptic compartments of R19G10 neurons. Bruchpilot (nc-82) staining (blue) visualizes neuropil. Above: brain imaged from anterior. Below: another brain, imaged from posterior. I & J) Mean speed of unexposed flies vs time for Clk4.1>TrpA1 and R19G10>TrpA1 genotypes and sibling controls, respectively. Shaded regions are +/- 1 standard error of the mean. Bars along x-axis indicate state of visible illumination (above) and temperature (below). K) Red light onset-triggered mean speed across flies of unexposed R19G10>CsChrimson flies vs time. -ATR indicates control flies not fed CsChrimson cofactor. Shaded regions are +/- 1 standard error of the mean. Bar along x-axis indicates lighting condition (black: darkness, red: red-light illumination).

Our manipulations targeted low-level biological elements (single genes and sparse neuronal expression patterns, as well as some broad expression patterns). To determine what higher level systems might be *E. muscae*’s target, we looked for enrichment of large effect sizes in the genes (or circuit elements) involved in the same higher level functions (or brain regions). We found that neurons in the antennal mechanosensory and motor center (AMMC), subesophageal ganglion (SOG), circadian system, and pars intercerebralis (PI) exhibited disproportionate abundance in top quintile of negative effect size (Fig 3D). Underscoring the potential importance of the PI, we observed that many of the neurons of large effect in the AMMC, SOG, and circadian system also innervated the PI. A comparable analysis for our genetic manipulations showed a clear enrichment for genes expressed in circadian cells (Fig 2E). Thus, our screen pointed conspicuously toward roles for the PI and the circadian network in summiting behavior.

**Table 2.**
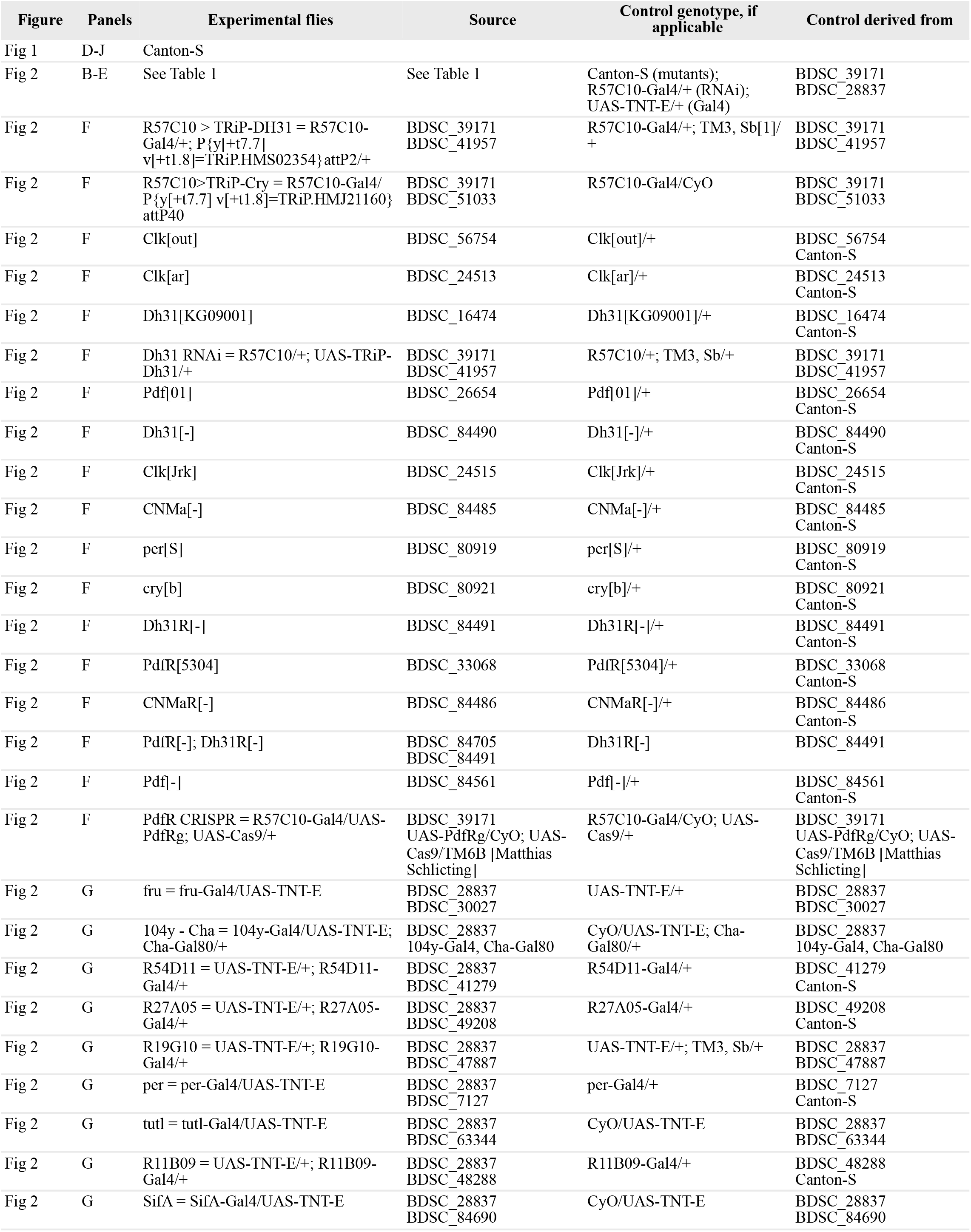
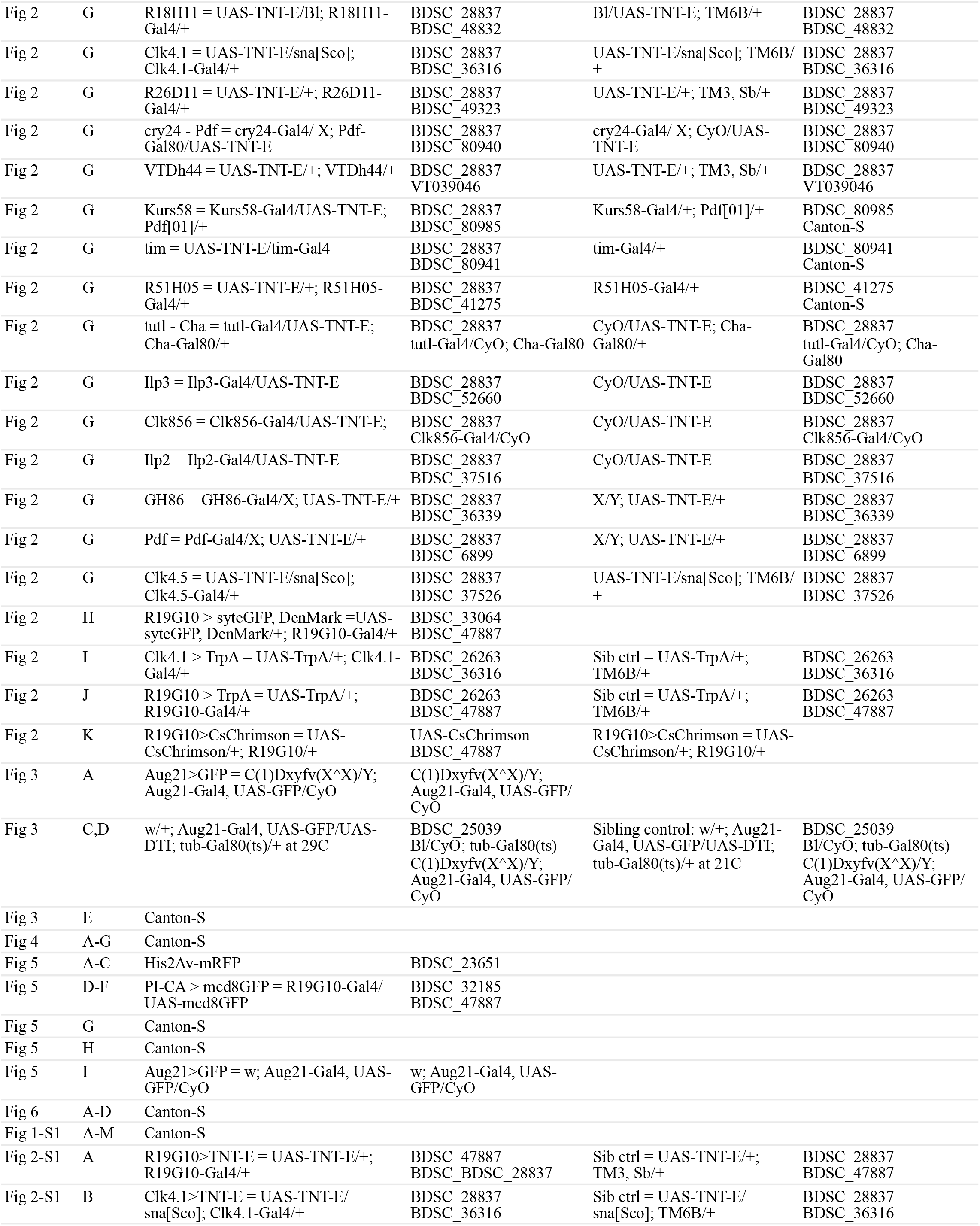
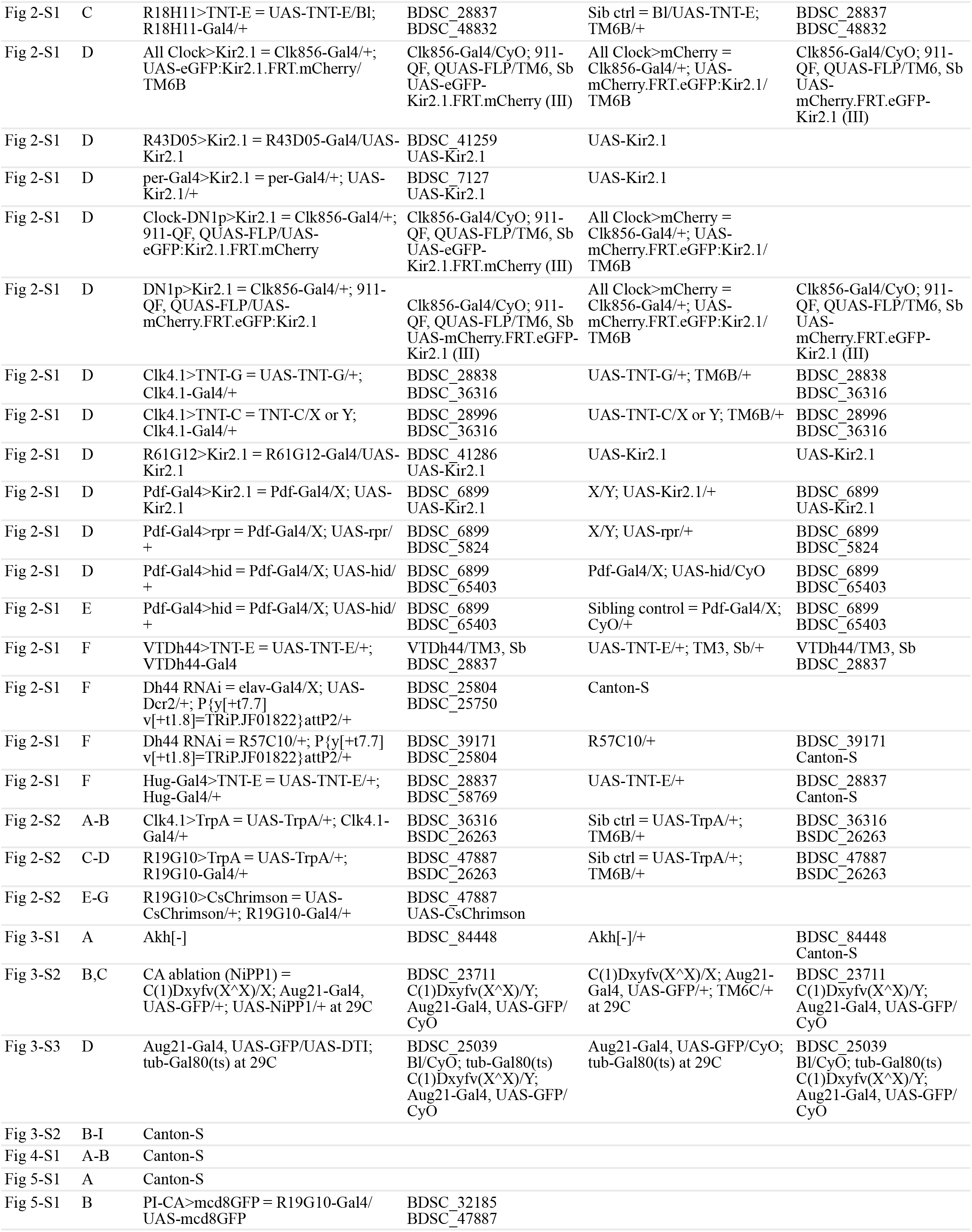
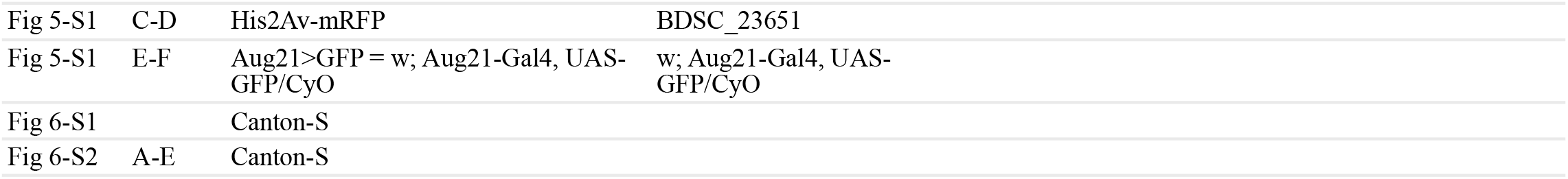
Genotypes of flies in manuscript figures and their paired control genotypes.

**Figure 3.**
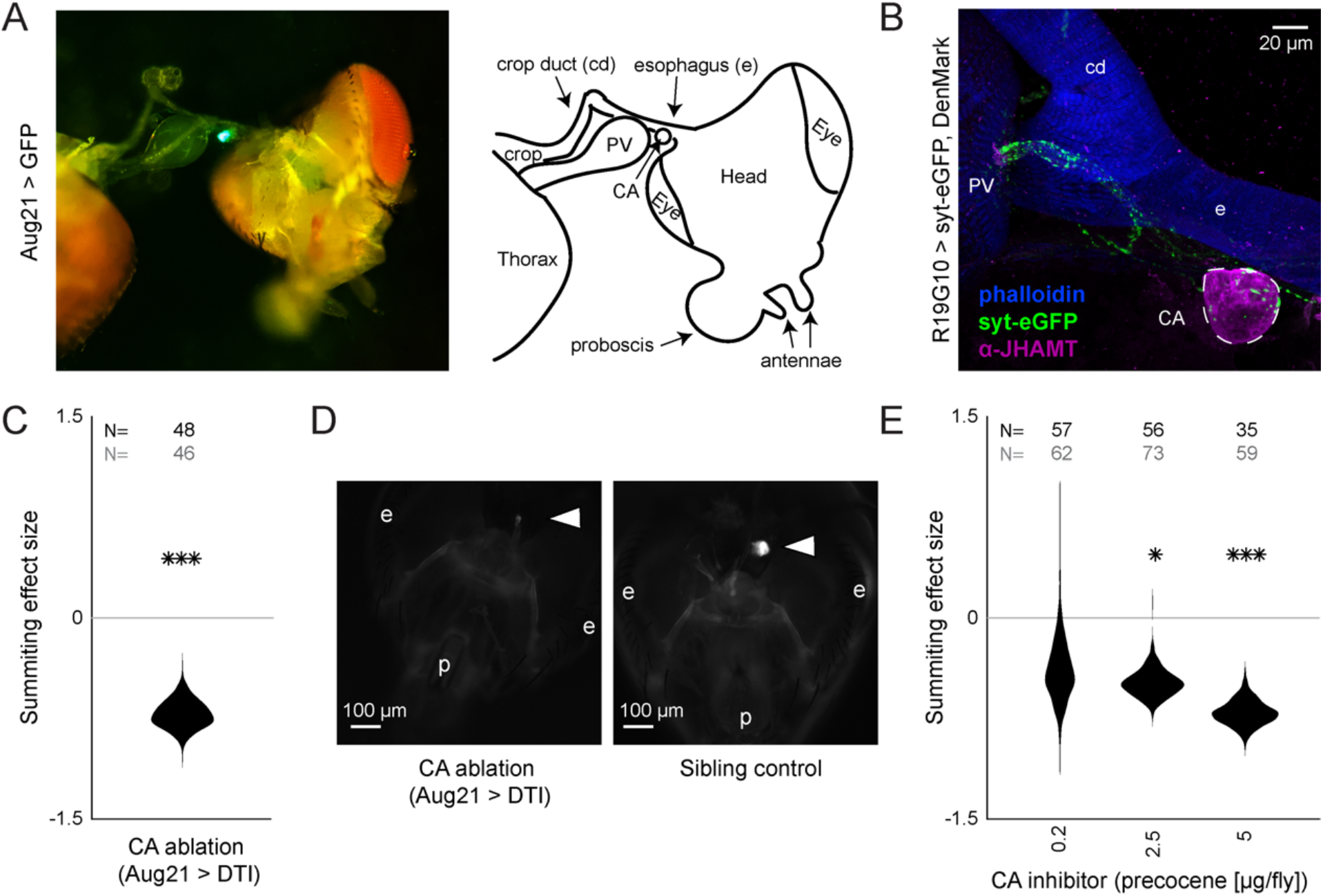
R19G10 (PI-CA) neurons project to the corpora allata, which are required for summiting behavior. A) Left: Composite micrograph of dissected Aug21>GFP fly, showing GFP fluorescence in the corpora allata (CA) overlaid on brightfield image. Right: Diagram of A with anatomical features labeled. PV = proventriculus. B) Confocal micrograph of immunostained RC from an R19G10>syt-eGFP, DenMark fly. Synaptic terminals are visible as green puncta, including in the CA. Magenta is anti-JHAMT and marks the CA. Blue phalloidin counterstain marks actin. Labels as in A. C) Summiting effect size estimate distribution of ablating the CA with diptheria toxin (DTI). Effect size is calculated relative to effector-less sibling controls. D) Micrographs of CA-ablated and effector-less, sibling, temperature-matched control flies (additional examples in Fig 3-S1D). White arrows indicate expected location of CA. e = eye, p = proboscis. E) Summiting effect size estimate distributions of various concentrations of the CA-ablating drug precocene. Effect size is calculated relative to vehicle (acetone) control. For C & E, effect sizes were estimated as in Fig 2; asterisks indicate statistically significant effects (* = p<0.05; ** = p<0.01; *** p<0.001). Sample sizes of experimental and control experiments are given in black and gray, respectively.

With these high level systems implicated as targets of fungal manipulation, we returned to a granular analysis to determine what specific circuit elements in circadian cells and the PI best recapitulated the high level effects. We measured the summiting response of an individually tailored genetic control for each circadian gene and PI or circadian circuit element (rather than screen-wide controls), and recalculated the effect size of each perturbation (Fig 2F,G). With respect to the circadian experiments, eleven mutants (Fig 2F) and four Gal4 lines (Fig 2G) showed impaired summiting compared to matched genetic background and/or sibling controls. We noticed that a large fraction of these hits affected a subtype of clock neurons, the group 1 posterior dorsal neurons (DN1ps). Silencing neurons with two DN1p drivers (Clk4.1 and R18H11; Kunst et al., 2014; Zhang et al., 2010) via TNT-E expression reduced summiting by 24-25% (p = 0.005, 0.019; Fig 2G, Fig 2-S1B,C). However, silencing the entire population of DN1p neurons by driving the inward-rectifying potassium channel Kir2.1 (Baines et al., 2001) with a pan-DN1p driver had no apparent effect (Fig 2-S1D). Genetic disruption of two signaling molecules expressed in DN1ps, Diuretic Hormone 31 (*Dh31*) and the neuropeptide CNMamide (*CNMa*), reduced summiting by 59 to 72% (3e-16 < p < 0.025; Fig 2F). Taken together, these results implicate DN1ps as mediating fungal manipulation while also revealing fine-scale complexity, as activity in some DN1ps, but not others, is required for full summiting.

DN1ps receive inputs from pacemaker neurons called small ventrolateral neurons (sLNvs) (Kaneko and Hall, 2000) that express the neuropeptide Pigment-dispersing factor (Pdf Helfrich-Förster and Homberg, 1993; Renn et al., 1999). While one *Pdf* mutant (*Pdf01*) exhibited a significant reduction in summiting (67%; p = 1.8e-16; Fig 2F), we saw no effect with another mutant whose *Pdf* locus was completely replaced (*Pdf-*). Disrupting sLNVs by expressing TNT-E, channel Kir2.1, or pro-apoptotic protein hid (Grether et al., 1995) also had no effect on summiting (Fig 2-S1D, E). This suggests that the main population of clock neurons upstream of DN1ps is irrelevant for summiting.

DN1ps send their axons medially, with many projections terminating at or near the PI (Kaneko and Hall, 2000). We tested the effect on summiting of silencing neurons in the PI using 16 different Gal4 drivers. Of these, seven produced significant reductions in summiting ranging from 44 to 79% (2.6e-9 < p < 0.02; Fig 2G). While some of these drivers were quite broad (such as *fru-Gal4*), others were quite sparse and specific to the PI, including *R19G10-Gal4* which is expressed in ∼12 neurons (all but two of which are in the PI; Fig 2H). Silencing R19G10 neurons reduced summiting by 60% (p = 2.4e-8; Fig 2G, Fig 2-S1A). Given the sparseness of this Gal4 driver and the large effect on summiting of expressing TNT-E with it, we focused on its PI neurons as the likely target of manipulation in this neuropil.

We next tested whether the ectopic activation of DN1ps or R19G10 neurons could drive “summiting” in flies that had never been exposed to *E. muscae*. We expressed a thermosensitive cation channel TrpA1 (Hamada et al., 2008) using *Clk4*.*1-Gal4* (to target DN1ps) or *R19G10-Gal4* (to target the PI) in flies unexposed to *E. muscae*. We conducted a 20h summiting assay with these flies, raising the temperature from 22°C to 28°C, for 2h (ZT6-8) between the flies’ daily circadian activity peaks that occur at the light-dark transitions (ZT0 and ZT12). Activating either DN1p or R19G10 neurons in this way led to an 28.7-fold or 9.7-fold increase in mean fly speed compared to sibling controls, respectively (Fig 2I and Fig 2J). This effect was significant across both males and females, though the effect was smaller in females for both experiments (Fig 2-S2A, S1B, S1C, S1D). As another test of the sufficiency of activating R19G10 neurons to induce summiting-like behavior, we expressed the optogenetic reagent CsChrimson (Klapoetke et al., 2014) in these cells. We ran these flies in a modified summiting assay with alternating periods of 3 minutes of darkness and red light. R19G10>CsChrimson flies fed all trans retinal (ATR), the CsChrimson cofactor, exhibited a burst of mean speed for the first 60s after light onset (Fig 2K, Fig 2-S2G) and suppressed walking speed for the last 90s of light stimulation, perhaps due to depolarization block (Herman et al., 2014). In contrast, control fly speed remained roughly constant throughout. The higher mean walking speed reflects a higher portion of flies walking after light onset (Fig 2-S2E,F). Thus, ectopically activating DN1Ps and R19G10 neurons appears to robustly induce a summiting-like increase in activity in flies unexposed to the fungus.

### The corpora allata are post-synaptic to R19G10 (PI-CA) neurons and necessary for summiting

In insects, pars intercerebralis neurons often project to the neurohemal organs of the retrocerebral complex (RC) (Carrow et al., 1984; de Velasco et al., 2007; Hartenstein, 2006; Pipa, 1978; Rüegg et al., 1983; Siegmund and Korge, 2001). We suspected this might be the case for R19G10 neurons. The RC in *Drosophila* consists of two pairs of fused neurohemal organs: the corpora cardiaca (CC) and the corpora allata (CA) (Nässel, 2002), the sole sites of adipokinetic hormone (Akh) (Noyes et al., 1995) and juvenile hormone (JH) synthesis, respectively (Klowden, 2008). Akh null mutants exhibited intact summiting (Fig 3-S1A), so we focused on potential R19G10 connections to the CA. We expressed the presynaptic marker synaptotagmin-GFP in R19G10 neurons and co-stained dissected brain-RC complexes for the CA-specific marker JH methyltransferase (JHMAT) (Niwa et al., 2008) (Fig 3-S2A). We observed R19G10 presynaptic terminals at the CA (Fig 3B), so we named R19G10 neurons “PI-CA” neurons to reflect this connectivity (Following the convention of Wolff and Rubin, 2018, the letters before the dash indicate the postsynaptic compartment, the letters after the presynaptic compartment).

To test if the CA were required for summiting, we turned to genetic ablation. First, we drove the expression of Nuclear inhibitor of Protein Phosphatase type 1 (NiPP1) with a driver that targets the CA (Aug21; Siegmund and Korge, 2001). NiPP1 overexpression causes cell-autonomous lethality in a variety of cell types (Parker et al., 2002) and has been previously used to ablate the CA in adult flies (Yamamoto et al., 2013). Aug21>NiPP1 animals showed reduced summiting by 60% (p = 2.7e-5) (Fig 3-S1B), but immunohistochemistry showed that the degree of CA ablation varied by animal (Fig 3-S1C). In a second ablation approach, we used a temperature-sensitive Gal80 (McGuire et al., 2004) to repress the expression of diphtheria toxin (DTI) driven by Aug21 until flies had reached wandering 3rd instar (Bilen et al., 2013). Tub-Gal80(ts), Aug21>DTI flies housed at the restrictive temperature also showed reduced summiting (72% (p = 1.1e-5, Fig 3C) and were confirmed by microscopy to have either greatly reduced or absent CA (Fig 3D, Fig 3-S1D).

We used pharmacology as a complementary approach to confirm the role of the CA in summiting. First, we blocked the production of JH by feeding flies fluvastatin, a compound that targets the JH synthesis pathway by inhibiting 3-hydroxy-3-methylglutaryl coenzyme A (HMG-coA) (Fig 3-S2A) (Debernard et al., 1994). Flies fed with fluvastatin at 72 hours after exposure to the fungus showed severely reduced summiting (110% (p = 3.1e-11) Fig 3-S2B). However, these flies released very few spores compared to untreated zombies and died at atypical times (after sunset; Fig 3-S2C). This observation led us to suspect that fluvastatin was impairing fungal growth. A series of experiments confirmed that feeding fluvastatin to flies well in advance of summiting (24 h post-exposure) led to premature death of infected flies (Fig 3-S2D) and abolished the circadian timing of death (Fig 3-S2E). Altogether, these data indicate that while fluvastatin disrupted summiting, that effect was likely due to disruption of fungal growth. We next turned to precocene (Bowers, 1981), a natural product that reduces JH titers per Amsalem et al., 2014 by inducing CA necrosis (Pratt et al., 1980). Applying 2.5 or 5 µg of precocene to exposed flies led to 47% and 70% reduction of summiting behavior (p = 0.001 and 6e-6, respectively) (Fig 3D). Increased doses of precocene led to more off-target deaths in both exposed and control flies, suggesting that precocene toxicity is fungus-independent (Fig 3-S2F). Precocene treatment did not alter timing of death by *E. muscae* (Fig 3-S2G).

We wondered if we could enhance summiting by dosing flies with the juvenile hormone analog (JHA) methoprene (Cerf and Georghiou, 1972). We topically applied methoprene at two different concentrations (2.5 and 5 µg). Surprisingly, these treatments led to a statistically non-significant reduction of summiting by 22.2 and 30.9% (p = 0.13, 0.09, respectively; Fig 3-S2H). We also tried to rescue the effects of precocene, either by coapplication of methoprene (2.5 µg) or by feeding flies another JHA, pyriproxyfen (5 µg) (Riddiford and Ashburner, 1991). Neither of these treatments rescued the effects of precocene treatment (Fig 3-S2I). Overall, these results indicate that CA function is necessary for summiting, but that supplementing flies with JHA is not sufficient to elicit this behavior. It could be that the acute release of JH is critical for driving summiting or that the CA produces a specific cocktail of juvenile hormones that are not well mimicked by our drug treatments.

### A real-time, automated classifier for summiting behavior

There may be physiological and anatomical differences between summiting and non-summiting flies that reflect its causal mechanism. These correlates likely degrade by the time the fly dies, so real-time identification of summiting flies is needed. We developed an automated classifier to identify summiting flies and alert an experimenter real time. Our ground-truth data set for training the classifier was made from a data set of ∼20 h recordings of speed and y-position from 1,306 *E. muscae*-exposed Canton-S flies, 345 of which were zombies. Each of the zombie traces was manually annotated with the time of summiting onset and time of death. Based on these timepoints, every frame was labeled as “pre-summiting,” “during summiting” or “post-summiting.” Every frame from survivor flies was labeled as “never summiting” to reflect that they would not summit for the period of observation (Fig 4A).

**Figure 4.**
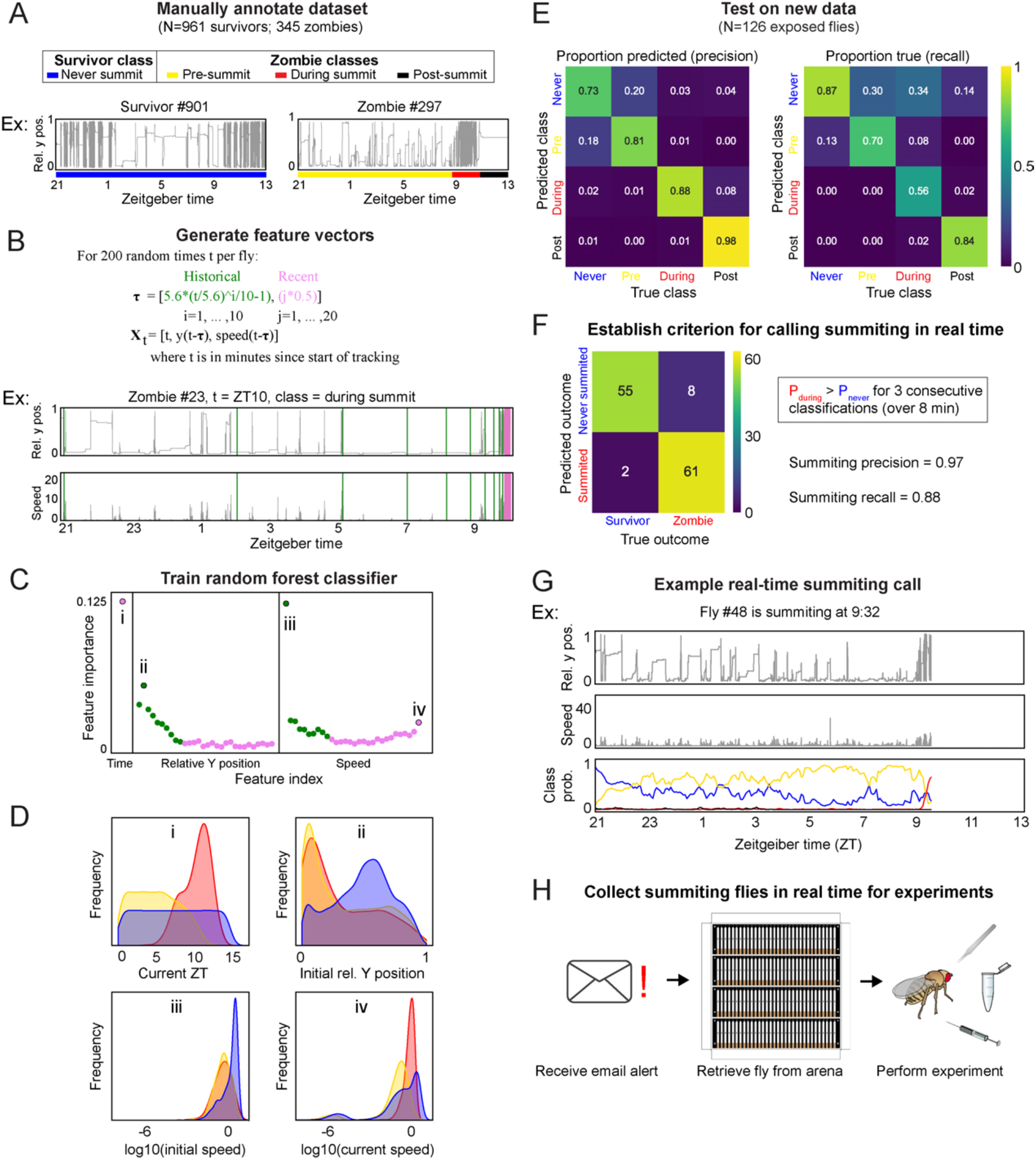
A random-forest classifier (RFC) for identifying summiting flies in real time. A) Top: classes learned by the classifier for zombies were pre-summiting = prior to onset of summiting (yellow), during summiting = after onset of summiting but before time of death (red), and post-summiting = after time of death (black). For survivors, there was one class, never-summiting (blue). Bottom: annotations of these classes on example y position trajectories from a survivor (left) and zombie (right). B) Feature vectors (Xt) generated for 200 random time points (t) for each fly. Vertical green and pink lines in example trajectory below indicate the historical (green) and recent (pink) values selected for the feature vector. C) Feature importance for classification of the 61 input variables. Roman numerals correspond to plots in subsequent panel. D) Distributions of important feature variables, visualized with kernel density estimation, across never summiting (blue), pre-summiting (yellow), and summiting (red) classes within the training dataset. E) Confusion matrices for precision (left) and recall (right) performance of the classifier on the test dataset. F) Confusion matrix for the survivor and zombie outcomes after implementing the real-time zombie-calling criterion. G) Example real-time behavior and class probability trajectories for a zombie fly, ending on the frame when it was called as a zombie. H) Summarized experimental workflow using the real-time classifier.

From each fly trajectory, we selected 200 random time points (for 261,200 total training data points) and from each generated a 61-element feature vector consisting of the current time, recent y-position and speed values, and past values of those measures log-spaced back to the start of the experiment (Fig 4B). Paired with each feature vector was the associated summiting label. We trained a random forest classifier with 75% of the data and validated performance with the remaining 25% (Fig 4SA). Of the variables in the feature vector, current time, initial y position, and initial and current speed were the most influential factors in classification (Fig 4C). The distributions of these variables by summiting label made sense: summiting labels were most abundant in the evening, at low y positions prior to summiting, and at higher speeds during summiting versus pre-summiting (Fig 4D). The classifier had middling recall (56%) but high precision (88%) on a novel test data set collected separately from the training and validation data (Fig 4E).

We next focused on how to use the classifier to flag summiting flies for upcoming real-time experiments. A rule wherein a fly was flagged as summiting when its during-summiting class probability exceeded its never-summiting class probability for three consecutive classifications (spanning 8 minutes) had high precision (97%) and recall (88%) (Fig 4F) in simulations of realtime experiments with ground truth labels (Fig 4G). Flies that never passed this threshold were flagged as “survivors”. Finally, we configured our fly-tracking software to run the classifier concurrently and email the experimenter when a summiting fly was flagged. Thus, we had a convenient, high-accuracy tool for experiments requiring real-time identification of summiting flies (Fig 4H).

### During summiting, E. muscae cells are adjacent to the PI and the PI-CA pathway appears intact

Using the real-time classifier, we assessed the distribution of *E. muscae* cells within the brains of summiting flies. We imaged the brains of summiting flies expressing RFP-tagged histones in all cells, counterstained with Hoechst to label all nuclei (fly and fungi). We observed a consistent pattern of *E. muscae* occupancy in the brain, with a plurality of fungal cells (27%-41%) in the superior medial protocerebrum (SMP), the region that contains the PI. Notably, there were very few fungal cells in the central complex, a premotor region (Fig 5A-C). Phalloidin staining suggested that each fungal cell sat in a “hole” in neuropil (Fig 5A). The dense occupancy of the SMP is established as early as 72 hours after exposure (Fig 5-S1A).

**Figure 5.**
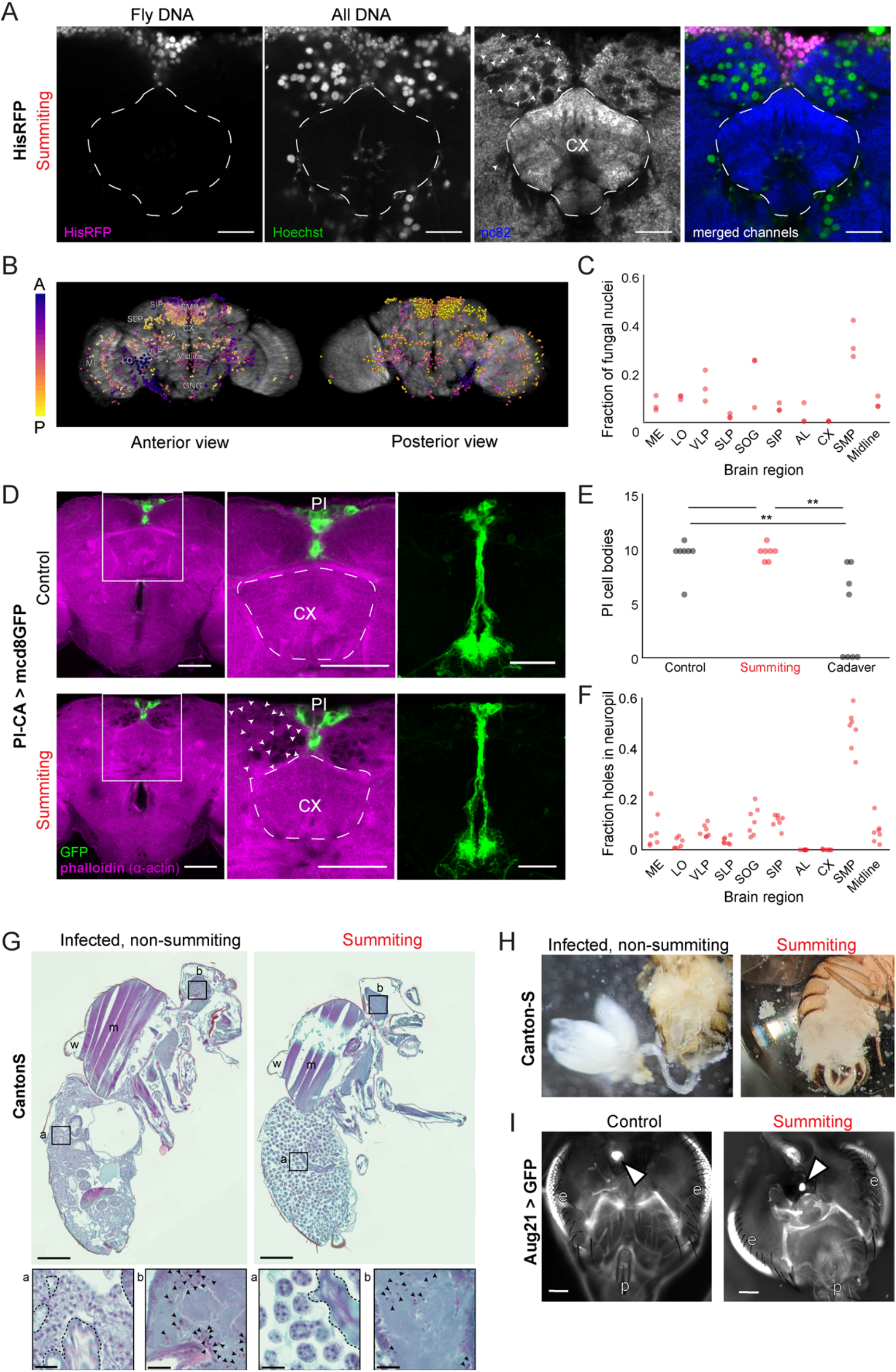
*E. muscae* densely occupies the SMP during summiting without apparent degradation of PI-CA neurons or CA. A) Confocal micrographs of the superior medial protocerebrum (SMP) from summiting His-RFP fly. Non-fly nuclei (Hoechst+, HisRFP-) are large compared to fly neuronal nuclei (Hoechst+, HisRFP+) and sit in “holes” in the neuropil visible in the nc82 counterstain channel. Scale bar is 20 microns. B) Whole brain invasion pattern of *E. muscae* (same brain as A). Nuclei are colored according to depth from anterior (A) to posterior (P). C) Distribution of fungal nuclei across brain regions (N=3). AL = antennal lobe, SIP = superior intermediate protocerebrum, SLP = superior lateral protocerebrum, CX = central complex, VLP = ventrolateral protocerebrum, SOG = subesophageal ganglion, LO = lobula, ME = medulla, midline = cells along midline of brain not in any other region. D) Confocal micrographs of PI-CA neurons (green) and phalloidin counterstain (magenta) in control and summiting flies. Left: sagittal planes of the central brain. Holes are apparent (in the phalloidin channel) in the SMP of the summiting brain, marked by arrowheads in one hemisphere. Holes are absent in CX of summiting brains and all control brain regions. Middle: Inset from left. Right: Maximum z-projections of GFP channel from full brain z-stacks. PI-CA morphology appears the same in summiting and control brains. Scale bars are 50 microns. E) Counts of PI-CA cell bodies in control (unexposed), summiting, or recently-killed (cadaver) PI-CA>mcd8GFP flies. F) Distribution of “holes” across brain regions. Abbreviations as in C. G) Safranin and fast green stained sections of paraffin-embedded Canton-S flies. Left: Infected, non-summiting fly (96 hours after exposure to fungus). Right: summiting, *E. muscae*-infected fly. a=abdomen, b=brain, w=wing, m=muscle. Scale bars are 200 microns. Insets of abdomen and brain are shown for each fly below (scale bars are 25 microns). Host tissues are outlined in dashed black; black arrowheads indicate fungal nuclei. H) Micrographs of dissected abdomens of 96 hour post-exposure non-summiting (left) and summiting (right) female flies. Gut and reproductive organs are still present in the non-summiting fly, but are absent in the summiting fly. Clumps of spherical fungal cells are visible in the dissection saline of summiting but not non-summiting fly. I) Fluorescence images of dissected Aug21>GFP flies. White arrowheads indicate CA. p=proboscis, e=eyes. Scale bars are 100 microns. Additional examples available in Fig 5-S1F.

To determine if the numerous *E. muscae* cells in the SMP were grossly disrupting PI-CA neurons, we imaged summiting animals expressing membrane-bound GFP in PI-CA neurons and compared them with uninfected controls. Despite the abundance of *E. muscae* cells in the SMP of summiting animals, the overall morphology of PI-CA neurons in summiting animals appeared normal (Fig 5D). There was no difference in the number of PI-CA cell bodies between summiting flies and unexposed controls (Fig 5E). In contrast, freshly killed cadavers had on average 60% fewer PI-adjacent cell bodies compared to summiting or non-summiting controls (0.0055 < p < 0.0029) (Fig 5E).

Fungal cells appear to displace host brain tissue, sitting in “holes” visible in actin-binding phalloidin counterstains (Fig 5A and 5D bottom middle). Consistently, the distribution of holes across brain regions (Fig 5F) was indistinguishable from the distribution of fungal nuclei (Fig 5C). Occasionally, we observed holes within the axon bundle of PI-CA neurons (Fig 5-S1B), but there was no indication of broken axons. Our interpretation is that during summiting, fungal cells displace neuropil without substantially consuming neural tissue or severing neural connections. This is consistent with the logic of zombie manipulation: *E. muscae* only consumes host tissues once they have served their purpose in aiding fungal dispersal.

While the brain is largely intact in summiting, this is not the case for organs in the abdomen, which are essentially obliterated in summiting flies (Fig 5G-H, Fig 5-S1E). The state of the abdominal organs is striking considering that these flies walk apparently normally. *E. muscae* in the abdomen of summiting flies adopted a spherical morphology distinct from their irregular protoplastic form before summiting, even as the interstices of the abdomen are packed with fungal cells (Fig 5G). *E. muscae* cells in the brain of summiting flies retain the appearance of pre-summiting hemolymph-bound cells (Fig 5G insets). The CA resides in the thorax adjacent to the esophagus and proventriculus. We wondered if these tissues might be degraded like the abdominal organs in summiting flies. We used the classifier to collect summiting and non-summiting Aug21>GFP animals, and found that the CA was consistently present in summiting flies (as well as controls) (Fig 5I, Fig 5-S1F). Overall, the preservation of the CA during summiting suggests that its function is needed to mediate summiting behavior.

### Evidence for the metabolic induction of summiting behavior

We wondered if *E. muscae*’s invasion of the brain disrupts the fly’s blood-brain barrier (BBB). Like vertebrates, flies maintain a BBB that restricts diffusion of compounds circulating in the hemolymph into nervous tissue (Hindle and Bainton, 2014). We assayed the integrity of the BBB of flies by injecting flies with Rhodamine B (RhoB), a fluorescent compound that is partially BBB-permeable (Pinsonneault et al., 2011). When RhoB enters the brain, it can be detected as fluorescence in the pseudopupil, the portion of eye ommatidia oriented toward the observer; high levels of RhoB can be observed as fluorescence across ommatidia (“bright eyes”) (Mayer et al., 2009). We found that BBB permeability was higher in exposed flies versus controls at 98 hours after exposure (Fig 6A). The increased permeability was not restricted to flies with confirmed infection (59% bright eyes), but was broadly observed among flies that had encountered the fungus (85% bright eyes), compared to unexposed controls (10% bright eyes) (Fig 6A). The proportion of bright eyed flies was lower at earlier time points following *E. muscae* exposure: 0% after 21 hours, 4.3% after 45 hours, 21.8% after 69 hours (Fig 6-S1). Our data are consistent with BBB permeability increasing with time since exposure.

**Figure 6.**
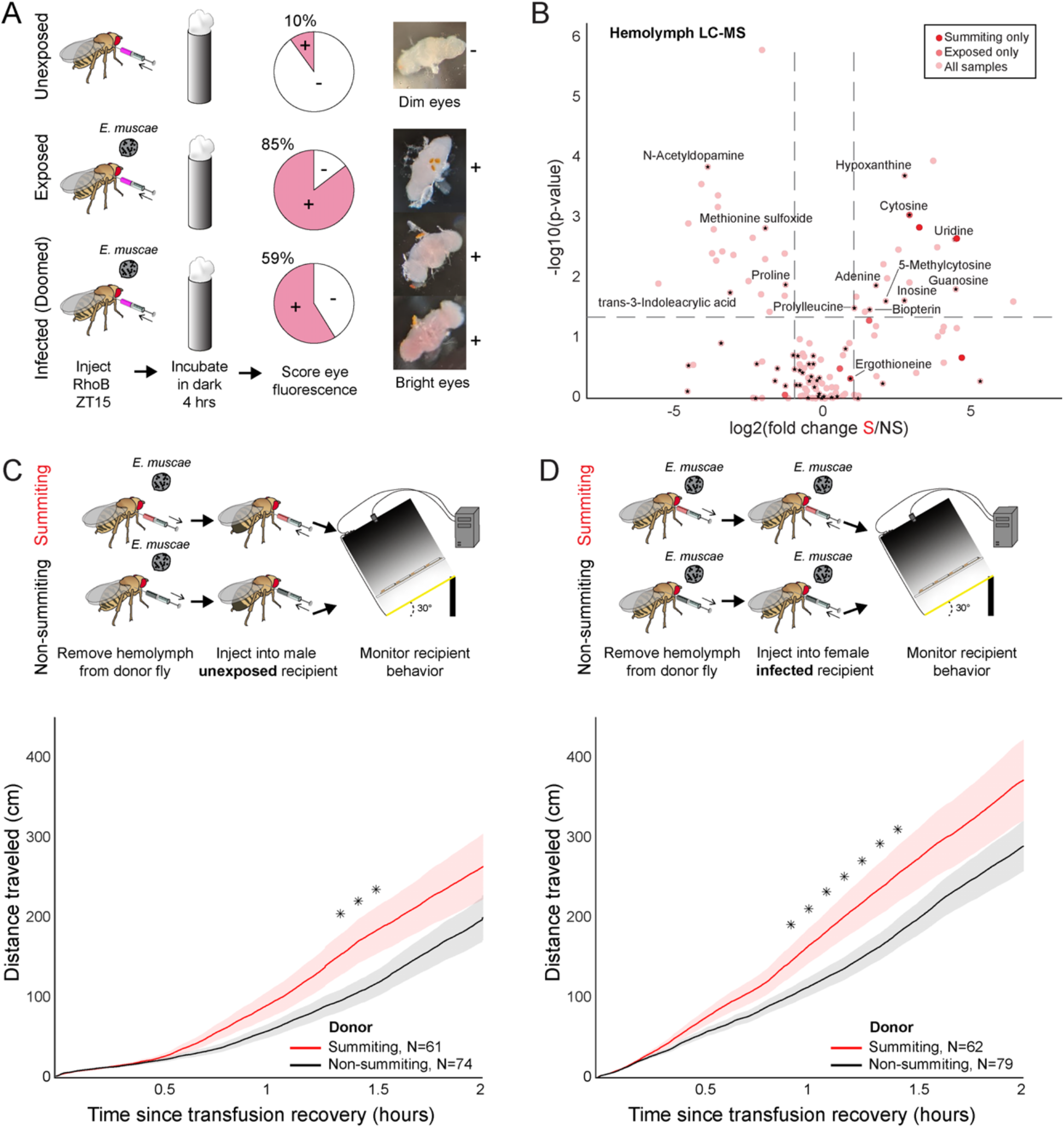
Hemolymph of summiting flies has a distinct metabolome and induces locomotion. A) BBB permeability of *E. muscae* exposed (96 hours) or unexposed flies assessed as the portion of flies with eye fluorescence after RhoB injection (N-40-50 per group). Infected (doomed) flies are exposed flies with fungal growth visible by eye through abdominal cuticle, all of whom would go on to summit within 22 hours. Bright-eyed flies (+) had visible RhoB uptake. Representative brains from dim and bright-eyed flies are shown at right. B) Volcano plot of hemolymph metabolites detected by LC-MS mass spectrometry in summiting (S) versus exposed, non-summiting (NS) flies. Putative identifications are given for selected compounds. See File S1 for compound abundances and statistical details. C & D) Total distance traveled versus time for flies receiving a transfusion of hemolymph from summiting donors. Diagrams at top indicate hemolymph transfusion experiment configuration. Shaded areas indicate + / - one standard error. Asterisks indicate p-values < 0.05 for two-tailed t-tests performed at each timepoint.

We next used metabolomics to compare the molecular composition of hemolymph in summiting flies to that of exposed, nonsummiting flies. We performed this experiment twice: once staging animals by hand based on flightlessness, which occurs during mid to late summiting (Fig 1B), and a second time using our automated classifier. We found that 168 compounds were detected in both of these experiments (Fig 6B, Fig 6-S2A-C), with nine compounds enriched and two compounds depleted in summiting versus exposed, non-summiting flies (Fig 6-S2A; see File S1 for specific fold-changes and p-values). Many of the compounds could not be identified. These included three compounds that were uniquely detected in summiting flies (C6H8N2O3, C14H16N6O7 and C12H19N2PS) (Fig 6B). Three additional compounds (molecular weights 276.08, 179.08 and 429.15 Da) were significantly greater in summiting versus exposed, non-summiting flies (6-S2A, File S1). Similarly, one compound of molecular weight 451.27 Da was significantly depleted in summiting flies (6-S2A, File S1).

Seventy-two compounds could be putatively identified. Cytosine was undetectable in the hemolymph of unexposed flies, but present in both exposed, non-summiting and summiting exposed flies (Fig 6B, Fig 6-S2A). Cytosine was significantly enriched in summiting versus exposed, non-summiting exposed flies (Fig 6B, Fig 6-S2A, File S1). Ergothioneine, an amino acid produced by some plants and microbes, including fungi (Borodina et al., 2020), was only detected in *E. muscae-*exposed animals (Fig 6-S2A), but did not appear to vary between summiting and exposed, non-summiting flies (Fig 6B). A handful of putatively identified compounds were present in all samples, but had significantly higher abundance in summiting flies versus exposed, non-summiting flies. These included uridine, guanosine and 5-methylcytosine (Fig 6B, Fig 6-S2A, File S1). Other putatively identified compounds were more abundant in exposed, non-summiting versus summiting flies: N-acetyldopamine, methionine sulfoxide and trans-3-Indoleacrylic acid (Fig 6B, Fig 6-S2B,C). Overall, these data indicate that summiting fly hemolymph is distinct from that of exposed, non-summiting flies.

To determine if factor(s) in the hemolymph of summiting flies could cause summiting behavior, we transfused hemolymph from summiting donors to non-summiting recipients, and tracked their ensuing behavior. We performed this experiment using exposed female donors and naive (unexposed) male recipients. Males tend to be smaller than females, so this choice of sexes maximized the quantity of hemolymph we could extract while minimizing its dilution in recipients. We observed a modest (37%) but significant increase in the distance traveled between 80 and 90 minutes post-transfusion, in flies that received summiting hemolymph compared to controls that received non-summiting hemolymph (0.033<p<0.039; Fig 6C). We conducted a second version of this experiment, this time with fungus-exposed females as the recipients and observed a similar increase in total distance traveled within the first 55-85 min after transfusion (44% increase, 0.024<p<0.048; Fig 6D). It is apparent that the hemolymph carries factors that can induce a summiting-like increase in locomotor activity.

### A neuro-mechanistic framework for summiting behavior

Altogether, our experiments point to a series of mechanisms by which *E. muscae* induces zombie summiting behavior (Fig 7). The fungus invades the brain as early as 48 hours prior to death (Elya et al., 2018), establishing extensive SMP occupancy by at least 24 hours before death. When summiting behavior begins ∼2.5 hours prior to death, the fungus has altered host hemolymph, likely via secretion of secondary metabolites. We hypothesize that these metabolites lead to the activation of PI-CA neurons, potentially via upstream DN1p clock neurons. In turn, we suspect that PI-CA activation stimulates the CA, leading to the release of JH. This hormone ultimately feeds back on the nervous system to generate the increase in locomotion at the heart of summiting. This framework unites the observations from many experiments and provides several specific hypotheses that we aim to tackle in future work.

**Figure 7.**
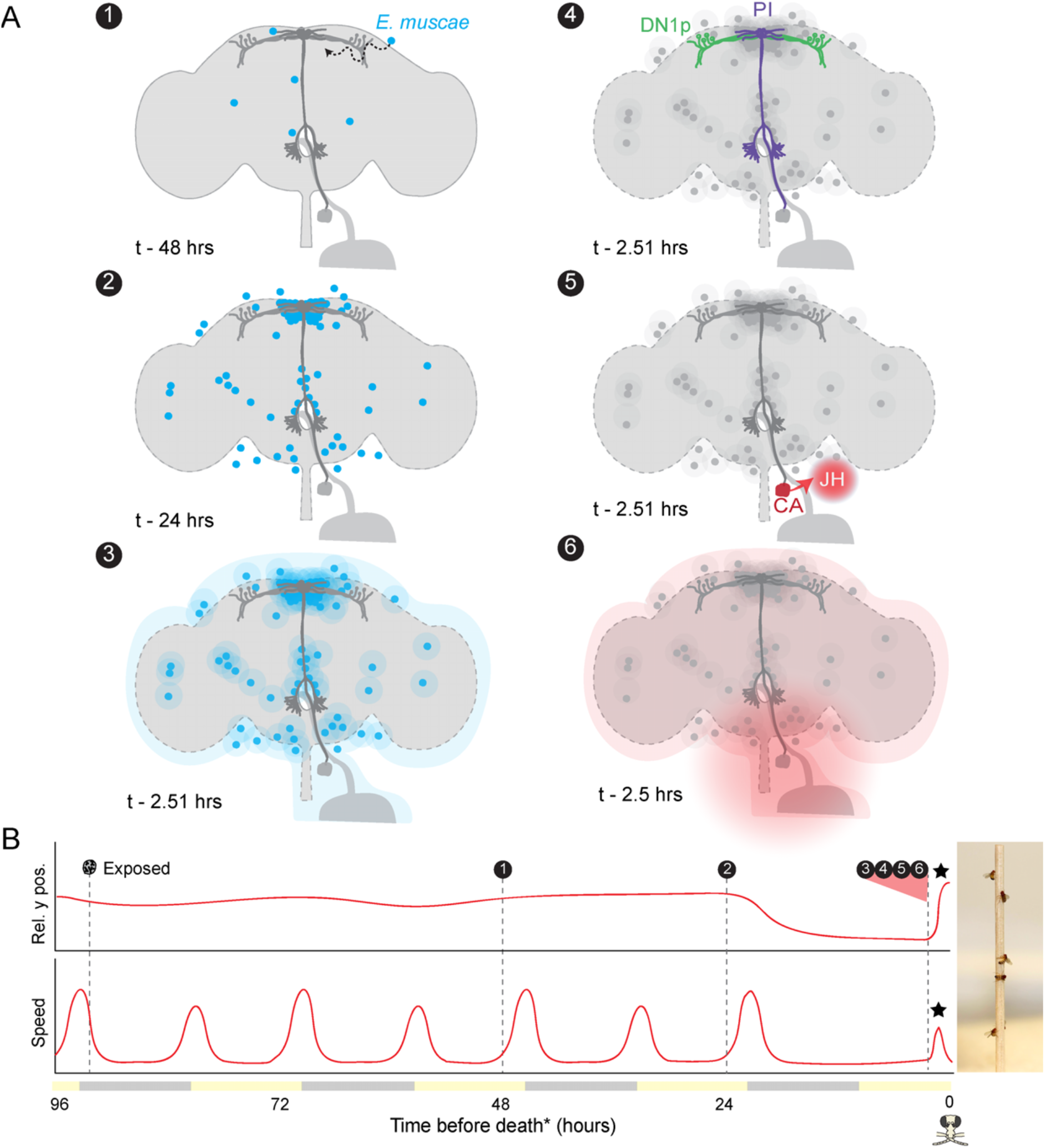
Proposed sequence of *E. muscae*-induced summiting mechanisms in zombie flies. A) Events in the host brain leading to *E. muscae*-induced summiting. (1) *E. muscae* cells are present in the brain as soon as 48 hours prior to death (Elya et al., 2018). (2) By 24 hours prior to death, the fungus is present at high density in the SMP. (3) *E. muscae* alters the hemolymph (perhaps by secreting compounds, as depicted here) to trigger the onset of summiting behavior. (4) Hemolymph-borne factors alter the activity of the circadian network/DN1p and PI-CA neurons. (5) JH is released from the CA following changes in PI-CA activity. (6) Increased JH levels drive an increase in locomotion. The dashed outline of the brain becomes more prominent between steps 1 and 3 to reflect an increase in BBB permeability over these timepoints. B) Left: Timeline of events depicted in (A) overlaid on cartoon plot of average relative y position (above) and speed (below) for zombie flies. Summiting is indicated by a black star; death (time of last movement) is indicated by a fly “skull.” Right: Zombie flies summited on a wooden dowel.

## Discussion

The discovery of dead, fungus-covered flies in elevated locales has fascinated the scientifically curious for at least the past 150 years (Berisford and Tsao, 1974; Cohn, 1855; Gryganskyi et al., 2013; Mullens, B A Rodriguez, J L Meyer, J A, 1987). Until very recently the biological mechanisms determining how they got there have been purely a matter of guesswork. Here, we reported a multi-pronged approach to characterize summiting behavior in zombified flies and make the first substantial progress towards understanding its mechanistic underpinnings using the *E. muscae-D. melanogaster* “zombie fly” system.

### A new understanding of summit disease

By analyzing the behavior of hundreds of *E. muscae*-exposed wild-type Canton-S flies in a custom summiting assay (Fig 1C), we discovered that a signature of summit disease is a burst of locomotor activity in the final ∼2.5 hours of a zombie fly’s life (Fig 1F-H). If the fly was previously in a low position, such as on the ground, or, in our assay, on the food, the net effect of increased activity will be upward motion. Perhaps it may be easier for parasites to evolve to manipulate neural mechanisms underlying activity in general, rather than the more specific circuits mediating negative gravitaxis. Notably, flies tend to die in higher positions when they begin summiting in the middle of a long arena (as determined by the positioning of the food) (Fig 1-S1I). This implies that *E. muscae* induces both increased activity and negative gravitaxis (to some degree), which interact with the geometry of the arena and the position of the fly prior to behavioral manipulation, to produce the summiting phenotype. Enhanced locomotor activity (ELA) is emerging as a recurring theme in insect behavior manipulation, having now been reported as a result of parasitism by not only fungi (Boyce et al., 2019; Trinh et al., 2021) but also viruses (Kamita et al., 2005; van Houte et al., 2012). It remains to be seen if other known examples of ELA are driven by similar mechanisms as by *E. muscae* and whether ELA is a universal feature of parasite-induced summit disease (e.g., in *Entomophaga grylli*-infected grasshoppers and *Pandora formica*(Małagocka et al., 2017) and *Dicrocoelium dendriticum*-infected ants; Pickford and Riegert, 1964; Martín-Vega et al., 2018).

### Host circadian and pars intercerebralis neurons mediate summiting

We leveraged our high throughput assay to screen for fly circuit elements mediating summiting and found evidence for the involvement of circadian and neurosecretory systems (Fig 2A-E). We identified two specific neuronal populations important for summiting: DN1p circadian neurons labeled by *Clk4*.*1-Gal4* (Fig 2F) and a small population of PI-CA neurons labeled by *R19G10-Gal4* (Fig 2G). Silencing these neurons significantly reduced summiting and ectopically activating them induced a summiting-like burst of locomotor activity (Fig 2I-K). These neurons are likely part of the same circuit; the projection of DN1ps to the PI has been confirmed both anatomically (Cavanaugh, et al., 2014) and functionally (Barber et al., 2021)

The pathway formed by these neurons is reminiscent of a previously characterized circadian-locomotor pathway. Cavanaugh et al (2014) showed that sLNv pacemaker neurons signal via DN1ps to a subset of PI neurons expressing the neuropeptide Dh44. Dh44-positive PI neurons project to a population of hug-in-positive neurons in the subesophageal ganglion (SOG), some of which send descending processes to the VNC (Cavanaugh et al., 2014; King et al., 2017). Recently, neurons that express both hugin and Dh44 receptor 2 (putatively the hugin*+* SOG neurons in (King et al., 2017)) were found to project to the CA (Mizuno et al., 2021). We did not observe a decrease in summiting by silencing or ablating sLNvs (Fig 2-S1D) or by silencing Dh44*+* PI neurons (Fig 2-S1F). However, we did observe an effect of silencing hugin*+* neurons (Fig 2-S1F). While it remains to be seen if any PI-CA neurons express Dh44, it is likely there are multiple connections between the PI and neurosecretory organs, and these pathways collectively exert control over locomotion. In the future, defining the neuropeptide profiles of PI-CA neurons may provide insight into the parasite’s proximate manipulation mechanism.

Silencing PI-CA neurons or mutating *Dh31* blocked summiting almost entirely, but silencing DN1p neurons had an effect that was roughly half as large (Fig 2G). This could reflect heterogeneity of DN1p cells (Ma et al., 2021). Another possibility is that additional inputs to PI-CA also mediate summiting manipulation, perhaps the Lateral Posterior clock Neurons (LPNs), which were also recently discovered to express Dh31 (Reinhard et al., 2021). The evolutionary logic of targeting the circadian network is elegant: strains of *E. muscae* have been reported to infect and manipulate a diverse collection of dipteran hosts (Elya and De Fine Licht, 2021). The proximate motor circuits controlling locomotor activity may vary from species to species, but all flies have a clock (Helfrich-Förster et al., 2020; Sandrelli et al., 2008) and the clock exerts a strong influence on locomotor behavior. Targeting the clock network and downstream neurosecretory neurons may represent a simple, conserved mechanism to appropriately activate motor programs across host species.

### PI-CA neurons induce summiting via their connection to the corpora allata

A defining feature of PI-CA neurons is their expression of presynaptic markers at the CA (Fig 3B), the conserved sites of JH synthesis and release within insects. JH has been implicated in a variety of physiological and behavioral phenomena within insects broadly (Riddiford, 2020; Tsang et al., 2020) and within fruit flies specifically (Zhang et al., 2022). Importantly, JH is known to have sexually dimorphic effects (Belgacem and Martin, 2007; Wu et al., 2018). While thermogenetic activation of DN1ps and PI-CA neurons induced both males and females to locomote (Fig2-S2A-D), the effect was 22.4- and 6-fold stronger in males, respectively. This difference is consistent with previous work implicating JH and the PI in sexually dimorphic locomotion (Belgacem and Martin, 2002; Gatti et al., 2000) and supports our conclusion that the CA and JH are the major output of DN1p and PI-CA neurons with respect to summiting. Given the sexually dimorphic effects of JH and ectopic PI-CA activation, one might expect strong sexual dimorphism in zombie summiting, but this is not observed (Fig 1-S1M). We propose that the apparent absence of sexual dimorphism in summiting is a consequence of effective castration by the fungus. Histological data showed that summiting flies either have severely damaged gonads or lack them entirely (Fig 5G, I), similar to other instances of parasitic castration (Cooley et al., 2018; Ewen, 1966; Lafferty and Kuris, 2009). As JHRs are present in gonads (Abdou et al., 2011; Baumann et al., 2017), it follows that in the absence of these sexually dimorphic tissues, JH-mediated behavioral differences between the sexes would be minimized.

We showed that summiting was reduced in *E. muscae*-infected flies with ablated CA (Fig 3C) or when treated with the JH synthesis inhibitor precocene (Fig 3E). However, we did not observe exacerbated summiting behavior in animals that had been treated with the juvenile hormone analog (JHA) methoprene (Fig 3-S2H) or a restoration of summiting behavior when animals received JHAs in addition to precocene (Fig 3-S2I). We suspect that summiting is driven by an acute spike in JH starting ∼2.5 hours before death, and our JHA experiments did not have this timing: methoprene was delivered in a single burst 20 hours pri- or to summiting and pyriproxyfen was administered chronically via the food. Secondly, we have strong reason to believe that whatever we applied to the fly was also making its way to the fungus (recall that healthy flies treated with both fluvastatin and methoprene were fine, but that this treatment was lethal for exposed flies (Fig 3-S2D)). Thus, another possibility is that the fungus is metabolizing the JHAs before they have a behavioral effect.

The role of the CA in *E. muscae*-induced summiting is consistent with the growing list of examples of parasites exploiting host hormonal axes (Adamo and Robinson, 2012; Beckage, 2012; Herbison, 2017; Tong et al., 2021). The JH pathway, in particular, has been shown to be modulated by a variety of insect parasites, ranging from nematodes to baculoviruses (Ahmed et al., 2022; Jiao et al., 2022; Nakai et al., 2016; Palli et al., 2000; Saito et al., 2015; Subrahmanyam and Ramakrishnan, 1980; Sun et al., 2019; Zhang et al., 2015). While there is clear consensus that JH is involved in a multitude of host physiological and behavioral processes, the extent of JH’s activities in insects is still being uncovered. Our data reveal another role for JH in the fruit fly: mediating *E. muscae*-induced summiting behavior.

### Machine learning classification of summiting animals in realtime

Identifying the molecular and physiological correlates of summiting is challenging for several reasons: summiting behavior is subtle to a human observer, summiting lasts just a few hours within a specific circadian window, and flies’ small size makes procuring sufficient material non-trivial. To make such experiments possible, we developed an automated classifier to identify flies as early into summiting behavior as possible (Fig 4). The random forest algorithm (Breiman, 2001; Pedregosa et al., 2012) at the heart of our classifier identified time of day (evening), previous position (low), previous speed (low), and current speed (high) as key features identifying summiting flies (Fig 4 C,D). The classifier achieved excellent precision and good recall on a novel cohort of exposed flies. By interfacing the classifier with an email alert system, we created a robust, scalable pipeline for procuring summiting flies for a variety of downstream experiments (Fig 5 & 6B-D).

### Morphological correlates of summiting

Using our real-time classifier, we conducted a comparison of host morphology prior to and during summiting. Previous analyses of infection progression suggested that the fungus was not occupying the brain with any spatial specificity (Elya et al., 2018), but here we found otherwise. There is a clear pattern of fungal cells densely invading the SMP of summiting flies, a neuropil that harbors DN1p axons and PI-CA cell bodies and dendrites (Fig 5B,C,F). This concentration of fungal cells is apparent at least 72 hours after exposure to *E. muscae* (Fig 5-S1A). Fungal cells are present in the brain as early as 48 hours after exposure (Elya et al., 2018), and the exact timing of when they accumulate in the SMP remains to be established. The distribution of *E. muscae* across neuropils, which is consistent across animals (Fig 5C), is interesting both for where fungal cells are and are not found. Fungal cells are noticeably absent from the central complex, a pre-motor center (Bender et al., 2010; Strausfeld, 1976) that may be involved in coordinating walking during summiting. Though morphological examination suggested that fungal cells are displacing (Fig 5-S1B), rather than consuming, nervous tissue, more work is needed to determine if neurons are damaged or dying as a result of adjacent fungal cells.

We observed extensive degradation of host abdominal tissues in summiting animals (Fig 5G-H, Fig 5-S1E). We were stunned to find flies with obliterated guts and gonads walking apparently normally. Despite widespread destruction in the body, the CA and PI-CA neurons appear intact in summiting animals, which is consistent with an acute role in summiting. We speculate that the fungus might achieve preservation of these tissues by preferentially digesting remaining host tissues from posterior to anterior. However, just because PI-CA neurons and the CA are present doesn’t mean they are functioning normally or at all. Future work should assess the physiology of these cells throughout the course of *E. muscae* infection.

### Physiological correlates of summiting

We discovered that the permeability of the blood-brain barrier was increased in exposed flies, as determined by assaying RhoB retention in fly brains (Fig 6A, Fig 6-S1). Our data suggest that BBB integrity degrades by the end of infection (Fig 6-S1), rather than rapidly after fungal exposure (by 21 hours) or upon fungal invasion of the nervous system (around 45 hours). A variety of insults, including bacterial infection, can lead to increased BBB permeability in fruit flies (Kim et al., 2021). We speculate that the progressive reduction in BBB integrity may result from the growing burden of the infection as the flies become sicker and sicker. In addition, the permeability of the BBB fluctuates over the day in a clock-dependent manner (Zhang et al., 2018). If the host’s circadian system is disrupted during infection, this could also be a source of compromised BBB integrity.

We found that the hemolymph metabolome of exposed, summiting flies differs from that of exposed, non-summiting flies and healthy controls (Fig 6B, Fig 6-S1). Three compounds of putative chemical formulae C6H8N2O3, C14H16N6O7 and C12H19N2PS appeared unique to summiting flies but could not be identified further. These compounds are prime candidates for further studies. Seven other compounds were significantly more abundant in summiting versus non-summiting flies across our replicate experiments: three of these could not be identified (MW 276.08, 179.08 and 429.15 g/mol) and the other four were putatively identified as guanosine, uridine, cytosine and 5-methylcytosine. Future collection of large quantities of summiting flies and fractionation approaches could be used to home in on compounds of interest and determine their chemical structure such that these compounds can be produced synthetically and assayed for behavioral effects (Beckerson et al., 2022). Cytosine is a pyrimidine nucleobase used in both DNA and RNA, a core molecular building block. It is intriguing that it was only detected in fungus-exposed fly hemolymph. High levels of cytosine have also been detected in the hemolymph of *Beauveria bassiana*-infected silkworms (Xu et al., 2015) and the serum of Sars-Cov2-infected humans (Blasco et al., 2020), with cytosine levels actually being predictive of infection status. Notably, a major derivative of cytosine, 5-methylcytosine, is also more abundant in summiting than non-summiting hemolymph. We hypothesize that elevated levels of cytosine could be a general indicator of infection, and its specific correlation with summiting warrants further investigation.

We detected ergothioneine in flies exposed to the fungus, either summiting or non-summiting. Ergothioneine has been hypothesized to play a role in host tissue preservation in *Ophiocordyceps* manipulated ants (Loreto and Hughes, 2019). Our data are consistent with ergothioneine being produced by *E. muscae*, but are not consistent with ergothioneine being produced only during summiting.

We saw that N-acetyldopamine (NADA), methionine sulfoxide, and trans-3-indoleacrylic acid were more abundant in non-summiting versus summiting flies. NADA is a product of dopamine (DA) breakdown (Neckameyer and Leal, 2017) and has been found to inhibit CA synthesis of JHs in *Manduca sexta* larvae (Granger et al., 2000)). DA, on the other hand, has been detected in the CA of *Manduca sexta* (Krueger et al., 1990) and studies in bees suggest a positive correlation between dopamine (DA), JH, and activity (Akasaka et al., 2010; Mezawa et al., 2013).

To test whether hemolymph-circulating factors in summiting animals can cause an increase in locomotion, we transfused hemolymph from classifier-flagged summiting flies into fungus-exposed and non-exposed recipients (Fig 6C,D). In both of these experiments, recipient flies exhibited a significant increase in locomotion over ∼1.5 hours post-transfusion. The effect size was modest (40% increase in total distance traveled in that interval), but this was not surprising as 1) we could only extract and transfer very small quantities (MacMillan et al, 2014) of hemolymph between animals and 2) this small quantity was diluted throughout the whole recipient fly’s body. Overall, this experiment provides direct evidence that one or more factors in the hemolymph of summiting flies cause summiting.

### A mechanistic framework for summiting behavior and beyond

Our experiments have revealed key mechanisms likely to underlie the summiting behavior of zombie flies. *E. muscae* cells perturb the activity of circadian and neurosecretory neurons, leading to the release of JH and a resultant increase in locomotion. This effect is at least partially mediated by summiting-specific factors circulating in the hemolymph. Of course, many questions remain. What compounds mediate the effect of transfused hemolymph? What cells are targeted by these compounds and by what molecular mechanisms? Do the fungal cells need physical access to the brain to induce a full summiting response? Is the proximity of fungal cells adjacent to DN1p axons and PI during summiting merely a coincidence? Future work should use spatially-resolved transcriptomic, metabolomic, and immunohistochemical approaches to answer these questions.

It is likely there are yet-to-be-discovered circuit elements mediating summiting. Silencing PI-CA neurons or ablating the CA severely attenuated summiting, but did not completely eliminate it. The dispersal and survival of *E. muscae* depends on a robust summiting response in the host (Carruthers, 1981), and the coevolutionary relationship between these species likely extends back 200-400 million years (Boomsma et al., 2014; Elya and De Fine Licht, 2021). Such a robust strategy is unlikely to rely on a single perturbation that could be countered by simple evolutionary changes in the host. An increase in locomotion can be achieved in many ways and is the likely output of many different behavioral circuits (Bidaye et al., 2020; Cavanaugh et al., 2014; Lee et al., 2021), so it would be unsurprising to find that multiple host circuits are targeted, including others yet to be discovered. Nevertheless, our study has identified a host pathway that likely mediates the predominant effects of the zombie fly summiting manipulation. These discoveries were made possible by studying summiting in a genetic model organism using high throughput behavioral assays. These tools and more will be essential to answer the many exciting questions arising from this work.

## Methods

### Data availability

All data supporting these results and the analysis code are available at *http://lab.debivort.org/zombie-summiting/*. Raw behavioral tracking (centroid versus time) data (450 GB) are available by request via external drive.

### Fly stocks and husbandry

All fly stocks were maintained in vials on cornmeal-dextrose media (11% dextrose, 3% cornmeal, 2.3% yeast, 0.64% agar, 0.125% tegosept [w/v]) at 21°C and ∼40% humidity in Percival incubators under 12 hour light and 12 hour dark lighting conditions and kept free of mites. Fly stocks used for experiments are listed in Table 1; full genotype information by figure panel is given in Table 2. Imaging and metabolomic data is from female flies and behavior data comes from mixed-sex populations, unless otherwise specified in the text.

### E. muscae *husbandry*

A continuous *in vivo* culture of *E. muscae* ‘Berkeley’ (referred to herein as *E. muscae*) isolated from wild Drosophilids (Elya et al., 2018) was maintained in Canton-S flies cleared of *Wolbachia* bacteria following the protocol described in Elya et al., 2018 and summarized as follows. Canton-S flies were reared in bottles containing cornmeal-dextrose media (see Fly stocks and husbandry) at 21°C and ∼40% humidity under 12 hour light and 12 hour dark lighting conditions. *E. muscae*-killed flies were collected daily between ZT15 and ZT18 using CO2 anesthesia. To infect new Canton-S flies, 30 fresh cadavers were embedded head first in the lid of a 60 mm Petri dish filled with a minimal medium (autoclaved 5% sucrose, 1.5% agar prepared in milliQ-purified deionized water, aka “5AS”). Approximately 330 mg of 0–5-day-old Canton-S flies were transferred to a small embryo collection cage (Genesee #59-100) which was topped with the dish containing the cadavers. The cage was placed mesh-side down on a grate propped up on the sides (to permit airflow into the cage) within an insect rearing enclosure (Bugdorm #4F3030) and incubated at 21°C, ∼40% humidity on a 12:12 L:D cycle. After 24 hours, the cage was inverted and placed food-side down directly on the bottom of the insect enclosure. After 48 hours, the cadaver dish was removed from the cage and replaced with a new dish of 5AS without cadavers. Starting at 96 hours, the collection cage was checked daily for up to four days between ZT15 and ZT18 for *E. muscae*-killed flies. These were collected using CO2 anesthesia and used to infect additional flies for experiments as described below.

### Summit behavior box design and fabrication

The summit assay box was designed in Adobe Illustrator in the style of other high throughput behavioral assays used by our lab (See (Werkhoven et al., 2021); *https://github.com/de-Bivort-Lab/dblab-schematics*). Nine behavior boxes were assembled from laser-cut acrylic and extruded aluminum railing (80/20 LLC). Each box consists of a ⅛” black acrylic base supporting an edge-lit dual-channel white (5300K) and infrared (850 nm) light LED board (KNEMA), three ⅛” black acrylic sides, a ¼” black hinged door and a ⅛” black ceiling upon which is mounted a digital camera (ELP-USB130W01MT-FV) equipped with an 87C Wratten infrared longpass filter (B&H Video #KO87C33O). The summit arenas sit on a ⅛” clear acrylic board held 6-7 cm above the illuminator by fasteners in the aluminum rail supports. 850 nm infrared illumination (invisible to flies) is used for tracking and white illumination (visible to flies) provides 12 hour light:dark circadian cues. Intensity of infrared and white light were independently controlled by pulse-width modulation via a Teensy (v3.2) microcontroller mounted to a custom printed circuit board (PCB) (Werkhoven et al., 2019). Each box’s camera and PCB connect to a dedicated Lenovo mini-tower PC running Windows 10 and Matlab v.2018b equipped with MARGO v. 1.03, Matlab-based software optimized to track many objects simultaneously, to record centroid positions for each of the assayed flies (Werkhoven et al., 2019). A complete list of parts and instructions for fabricating a summiting box can be found at *https://github.com/de-Bivort-Lab/dblab-schematics*.

### Summiting behavior arena designs

Several different arena variants were used in the summiting assay tracking boxes. All arenas were fabricated in arrays in acrylic trays that fit snugly into the assay boxes. Each arena includes a small hole at one end through which a fly can be aspirated and subsequently sealed using a small cotton ball. Arenas were 3.2mm tall, allowing flies to walk freely and raise their wings, the final manipulation by *E. muscae*.

An early prototype summiting assay was angled at 90°, but we found that even with a sandpaper-roughened walking surface, dying flies struggled to maintain their grip on the vertical surface. This was manifested in two ways: 1) flies exhibited sudden, rapid downward movement in their behavioral traces consistent with falls and 2) *E. muscae*-killed flies were predominantly found at the bottom of the well at the end of the experiment. This was subsequently confirmed by reviewing video taken from these experiments. To remedy this, we reduced the incline to 30°, which is sufficient for flies to respond behaviorally to the direction of gravity (M. Reiser, personal communication). This eliminated obvious falling bouts and yielded a wide range of final positions ranging from the bottom to the top of the arena.

### Standard arena (e.g., Fig 1F)

Standard arenas measured 6.5 cm long by 0.5 cm wide by 0.32 cm tall and housed a single fly. Arenas were constructed in rows of 32 from three layers of ⅛” laser cut acrylic consisting of a clear base manually roughened with 120 grit sandpaper, black walls, and a clear top. The layers were held together with 8-32 screws and nuts. A 3 mm loading hole in the lid at one end of the arena permitted loading of an anesthetized fly with a paintbrush. This entry hole was sealed with a piece of dental cotton after the fly was loaded. A minimal medium, 5AS, was provided at the opposite end of the chamber. The end of the chamber with food was sealed with two layers of Parafilm to slow desiccation of the food. Fully prepared (i.e., with food at the bottom and the loading hole sealed), the long axis of the arena had ∼5 cm of open space. Each tray had four rows of arenas, for a total of 128 arenas per tray. Laser-cutting designs for the standard arenas are available at *https://github.com/de-Bivort-Lab/dblab-schematics/tree/master/Summit_Assay*.

### Starvation arena (e.g., Fig 1-S1A)

Starvation arenas were constructed as standard arenas, substituting 1.5% agar (no sucrose) for 5AS media.

### Desiccation arena (e.g., Fig 1-S1B)

Desiccation arenas were constructed as standard arenas, except each arena was 6 cm tall (∼5.7 cm effective height) and lacked food and any opening at the bottom for the introduction of food.

### Two-choice arena (e.g., Fig 1-S1F)

Two choice arenas consisted of a five-layer acrylic sandwich secured with 8-32 fasteners: a bottom layer consisted of a ⅛” clear base texturized with 120 grit sandpaper. The next two layers each consisted of 1/16” black walls dividing the row into 32 chambers. These layers were rotated 180° with respect to each other, leaving gaps in the floor and ceiling at opposite ends of the arena that could be filled with media. Thus, the total height of the arena, except at the ends, was 1/8”. Each chamber was 4.6 cm long and contained 5AS at one end, 1.5% agar at the other. The lid layer consisted of ⅛” clear acrylic. Flies were loaded quickly into the arenas and the lid was placed before the flies could wake up. Each tray had four rows of arenas, for a total of 128 arenas per tray.

### Tall arena (e.g., Fig 1-S1I)

Tall arenas were constructed in the same fashion as standard arenas but measured 13 cm high instead of 6.5 cm. Two rows of 30 tall arenas each filled each tray. Food was pipetted into the middle of each arena and allowed to cool before the arenas were inclined. Flies were loaded through a loading hole at one end of the arena. The hole was plugged with cotton, for an effective length of ∼12.8 cm.

### Summiting behavior experiments with E. muscae exposed flies

All summiting experiments with *E. muscae*-exposed flies were run as follows (unless otherwise indicated): flies of were exposed to *E. muscae* by first embedding eight sporulating Canton-S cadavers in a 2.3 cm-diameter disc of ∼3.5 mm thick 5AS that was transferred with 6” forceps into the bottom of an empty wide-mouth *Drosophila* vial (Genesee #32-118). A ruler was used to mark 1.5 cm above the top of the disc. 0-5 day old flies of the experimental genotype were anesthetized with CO2, and 35 (∼half male, ∼half female) were transferred into the vial. The vial was capped with a Droso-Plug (Genesee #59-201) which was pushed down into the vial until the bottom was level with the 1.5 cm mark. For each experimental tray, three vials of flies were prepared in this way to expose a total of 105 flies; one additional vial of 35 flies was prepared identically but omitted cadavers as a non-exposed control. Together, these four vials were sufficient to fill a tray of 128 arenas. All prepared vials were incubated in a humid chamber (a small tupperware lined with deionized water-wetted paper towels) at 21°C on a 12:12 L:D cycle. After 24 hours, the vials were removed from the humid chamber and the Droso-plugs were pulled to the top of the vial to reduce fly crowding.

After 48-72 hours in the incubator, flies were loaded into the arenas using CO2 anesthesia. Flies loaded into arenas during scotophase (dark period of their 12:12 L:D circadian cycle) were shielded from ambient light in a foil-lined cardboard box. To begin behavioral experiments, arena trays were placed in the summit assay box and flies were tracked starting between ZT17 and ZT20. Tracking proceeded until ZT13 the next day (day 4). If many flies remained alive, tracking continued until ZT13 the following day. Some experiments, particularly in periods of COVID-restricted lab access, ran unattended until ZT13 on day 6 or 7. This variation in the timing of the end of the experiment had no effect on our measured outcomes, since all behavioral data were analyzed with respect to times of fly death, and any tracking data after death were ignored.

Tracking data were collected at 3 Hz using the circadian experiment template (*https://github.com/de-Bivort-Lab/margo/tree/master/examples/Circadian*) in MARGO v1.03 (Werkhoven et al., 2019; *https://github.com/de-Bivort-Lab/margo*) with the following settings: white light intensity 50%, infrared between 70-100%, adjusted to provide the best contrast for tracking, tracking threshold = 18, minimum area = 10, min trace duration = 6. Default settings were used for other configuration parameters. After tracking concluded, flies were manually scored as either alive (coded as survival=1 and outcome=0), dead with evidence of E. muscae sporulation (survival=0, outcome=1), or dead with no E. muscae sporulation (survival=0, outcome=0). These annotations were saved in a metadata file accompanying each MARGO output file and used in downstream analyses.

### Summit behavior data analysis

For each tray of flies (N<=128), we generated an experiment metadata table that incorporated the manually-scored survival outcome described above as well as fly genotype, sex, and fungal exposure status (exposed or non-exposed). Experiment metadata along with tracking data were input into a Matlab-based analysis pipeline that proceeded through the following steps: 1) automatic denoising, 2) manual time of death calling, 3) behavioral trajectory alignment to time of death, 4) SM calculation, 5) effect size estimation. See *http://lab.debivort.org/zom-bie-summiting/*.

The automatic denoising algorithm scanned speed throughout the experiment and flagged any ROIs that exhibited more than 20 instances per day of experimental time greater than ∼40 mm/ s. This threshold was chosen based on examination of individual ROI speed traces as a value that would only be exceeded with noise. The bulk of noisy behavioral recordings arose when the flies’ position was erroneously tracked as moving along the long edges of the arenas. Denoising was achieved by reducing the horizontal width of the arena region-of-interest (ROI) and recalculating centroid trajectory until speed violations fell below threshold or the ROI was trimmed to nine pixels, at which point its data was discarded.

Time of death was called manually for every cadaver (N=∼23,500) by CE throughout this study by checking time-aligned plots of y position and speed. Time of death was estimated as the time the fly was last observed to exhibit walking behavior. Extremely slow changes in y-position and tracking jitter around a particular y-position were not considered to be walking behavior. These definitions were initially validated by comparing paired behavioral video and tracking data. ROIs were flagged if sparse tracking occurred or residual noise was so great that the time of death couldn’t be reasonably determined. These ROIs were dropped in subsequent analysis. For the gene and Gal4 screen (Fig 2B,C), scoring of time-of-death was not blind to fly genotype; for all subsequent experiments, times of death were scored blind to experimental group. Time of death was stored as a frame number in the experimental metadata file.

Denoised tracking data and experimental metadata with time-of-death calls were input into a script that performed the following tasks: 1) determined the earliest start time for all experiments and aligned all data relative to this timepoint. This was necessary as experiments were not all started at precisely the same time (e.g., one experiment may start at 5:08 pm, another at 5:24 pm); 2) categorized each fly-trajectory as either a zombie (cadaver), survivor (alive) or unexposed control (uninfected)), based on experimental metadata; 3) randomly assigned a “time of death” for survivor and control flies from the pool of observed times of death within cadavers for that genotype, to make data between groups more comparable; 4) align all fly behavioral (y position and speed) trajectories relative to their time of death; 5) output a variable containing aligned and original vectors of data by category (zombie, survivor, unexposed) for a given genotype.

To calculate the summit metric (SM) for each cadaver, we first determined the period of summiting. The beginning of summiting was defined as 2.5h before death. The speed trajectory was smoothed with a one hour sliding window average and the end of summiting was defined as the earliest moment when the smoothed speed dropped to the same level as the start of summiting. The speed trajectory was baseline corrected by subtracting the smoothed speed at the onset of summiting, and the area under the resulting curve during the period of summiting divided by the duration of summiting (end of summiting – start of summiting) was taken as the value of SM. Thus, SM has units of distance/time and is a measure of speed.

### Statistical tests

Summiting effect size estimate distributions were calculated by bootstrapping flies, separately in experimental and control groups, calculating the manipulation effect size as (mean(Experimental SM) – mean(Control SM))/mean(Control SM), over 1,000 resamplings. Distributions were plotted as kernel density estimates. Two-tailed unpaired t-tests were used to assess significance of differences between SM in experimental and control groups. All reported p-values are nominal. Confidence intervals on time-varying data were calculated by bootstrapping individual flies over 1,000 replicates and shading the original mean values and +/-1 standard deviation of the bootstrapped means.

### Thermogenetic activation of DN1p and PI-CA

Unexposed flies (up to 8 days post eclosion) were loaded into standard summiting arenas (5AS food placed at y position = 0, 30° incline) and were tracked starting at ∼ZT17 in a temperature-controlled room initially held at 21°C, below the activation temperature of TrpA (Hamada et al., 2008). At ZT5:30 the following day, the temperature setpoint of the environmental room was increased to 28°C. The room took approximately 30 minutes to reach the setpoint temperature. Temperature in the room was monitored via a Bluetooth Thermometer (Govee #H5075). At ZT7:30 (2 hours after the initial setpoint change), the setpoint was returned to 21°C. Flies were tracked until ∼ZT13, for a total tracking time of 20 hours. Temperature measurements taken concurrently with behavioral tracking were used to generate the heatmap strips in Figure 2I, J, etc.

### Optogenetic activation of PI-CA

Young (up to 3 days post eclosion), unexposed UAS-CsChrimson/+; R19G10-Gal4/+ flies were placed in narrow (24.8 mm diameter) foil-wrapped vials, in which either 10 µL of 100 mM all-trans-retinal (ATR; Sigma) in ethanol, a required cofactor for CsChrimson, or 10 µL of 70% ethanol had been applied to the surface of the food. Flies in both groups were transferred to freshly-applied ATR/ethanol vials every 2 days. After 8 days, flies were tracked in individual, circular 28 mm diameter arenas (Werkhoven et al., 2021) using MARGO under IR illumination. For Fig 2K, Fig 2-S2G, flies were tracked for 30 minutes. After 15 minutes of tracking in darkness, constant red light (3.15 µW/ mm2) was projected onto the behavioral arenas using an over-head mounted modified DLP projector (Werkhoven et al., 2019). For Fig 2-S2E,F, red light was delivered in 5 ms pulses at 5 Hz for 30 seconds using the same projector under the control of the MATLAB PsychToolBox package (*http://psychtoolbox.org/*). Each 30 second pulsed red light trial was followed by 65 seconds of darkness (the projector light path was manually blocked with black acrylic during these periods), for 38 trials, totaling one hour of tracking.

### Immunohistochemistry

Tissues (brains, ventral nerve cords and/or anterior foreguts with retrocerebral complexes) were dissected in 1x PBS from female flies and stained generally following the Janelia FlyLight protocol (Janelia FlyLight Team, 2015) as follows. Fixation, incubations and washes all took place under gentle orbital shaking. Tissues were fixed in 2% paraformaldehyde for 55 minutes at room temperature in 2 mL Protein LoBind tubes (Eppendorf 022431064). Fixative was removed and tissues were washed 4x 10 minutes with 1.5 mL PBS with 0.5% Triton X-100 (PBT). Tissues were then blocked for 1.5 hours at room temperature in 200 µL of PBT with 5% normal goat serum (NGS) before adding primary antibodies prepared at the indicated dilutions in PBT with 5% NGS (Table 3). Tissues were incubated with primary antibodies for up to 4 hours at room temperature then placed at 4°C for at least 36 hours and no more than 108 hours. Primary antibody solution was removed and samples were washed at room temperature at 3x 30 minutes in 1.5 mL PBT. Tissues were then incubated in 200 µL of PBT containing 5% NGS and secondary antibodies (Table 3) for two to four hours at room temperature before moving to 4°C for approximately 60 hours. Secondary antibody solution was removed and tissues were washed 3x 30 minutes in PBT. Samples were then mounted in a drop of Vectashield placed within one or more 3-ring binder reinforcer stickers, which served as a coverslip bridge. Slides were sealed with nail polish and stored in the dark at 4°C until imaging on an LSM 700 confocal microscope (Zeiss).

**Table 3.** Immunohistochemistry reagents.

| Target (antibody) | Fluorophore | Dilution | Manufacturer |
| --- | --- | --- | --- |
| Nc82 (mouse IgG1) | None | 1:40 | Iowa Developmental Studies Hybridoma Bank |
| dsRed (rabbit) | None | 1:250 | Takara Bio #632496 |
| GFP (chicken IgY) | None | 1:4000 | AvesLabs #GFP-1020 |
| JHAMT (guinea pig IgG) | None | 1:1000 | Ryusuke Niwa (Niwa et al., 2008) |
| Mouse IgMμ (goat ) | Cy5 | 1:400 | Sigma #AP500S |
| Rabbit IgG (goat) | AF568 | 1:250 | ThermoFisher #A11011 |
| Chicken IgY (goat) | AF488 | 1:800 | ThermoFisher #A11039 |
| Guinea pig IgG (goat) | AF568 | 1:400 | ThermoFisher #A11075 |
| Actin (phalloidin) | AF568 | 1:400 | ThermoFisher #A12380 |
| DNA | Hoechst 33342 | 1:1000 | ThermoFisher #H3570 |

### Genetic ablation of CA

CA of adult flies were completely or partially ablated following the methods of (Bilen et al., 2013) and (Yamamoto et al., 2013), respectively. For complete ablation (Fig 3C,D, Fig 3-S1D), virgin females of genotype *Aug21-Gal4,UAS-GFP/CyO* were crossed to males of genotype *UAS-DTI/CyO; tub-Gal80ts/TM6B* and reared at 21°C until progeny reached third wandering instar. At this point, progeny were either transferred to 29°C until eclosion or kept at 21°C. Progeny of the genotype *Aug21-Gal4,UAS-GFP/UAS-DTI; tub-Gal80ts/+* were then exposed to *E. muscae* and run in the summit behavior assay. In separate experiments to assess ablation efficiency, experimental and control female flies were dissected and examined using a compound epifluorescence microscope (Nikon 80i).

For partial CA ablation (Fig 3-S1B,C), virgin females of genotype *C(1)Dxyfv(X^X)/Y;Aug21-Gal4, UAS-GFP/CyO* were crossed to *UAS-NiPP1* males at 29°C. Experimental flies (*C(1)Dxyfv(X^X)/Y; Aug21-Gal4,UAS-GFP/+;UAS-NiPP1/+*) and sibling controls (*C(1)Dxyfv(X^X)/Y;Aug21-Gal4, UAS-GFP/ +;TM6C/+*) were exposed to *E. muscae* and run in the summit behavior assay. To assess ablation efficiency, experimental and control female flies were subjected to immunohistochemistry using anti-GFP and anti-nc82 primary antibodies and imaged on a LSM 700 confocal microscope (Zeiss).

### Pharmacological perturbation of CA

Precocene I (Sigma# 195855) and methoprene (Sigma# 33375) were diluted in acetone (Sigma# 179124) and applied topically to the ventral abdomen of CO2-anesthetized flies that had been exposed to *E. muscae* (72 hours prior) or mock unexposed controls. 0.2 µL of the compounds were applied per fly using a 10 µL Hamilton syringe (Hamilton #80075) with a repeater attachment (Hamilton #83700). Acetone-only flies served as a vehicle control. To avoid compounds cross-contaminating flies, anesthetized flies were placed on top of two layers of fresh filter paper and handled with a reagent-dedicated paint brush as soon as they had been dosed with the desired compound. The syringe was thoroughly flushed with acetone between compounds.

Solutions of pyriproxyfen (Sigma# 34174, dissolved in ethanol) and fluvastatin (Sigma# PHR1620, dissolved in ultrapure water) were individually pipetted onto the media in standard summit arenas prepared with 5AS in 5 µL volumes using a 250 µL Hamilton syringe (Hamilton #81101) and repeater attachment. Five µL of either ethanol or water were applied to a second set of arena media to serve as vehicle controls for pyriproxyfen and fluvastatin, respectively. Arenas were then parafilm-sealed and stored at 4°C overnight. The following day, chambers were allowed to warm to room temperature before introducing flies for summit behavior assays.

### Real-time summiting classifier

A ground truth dataset was pooled from 14 experiments comprising 1306 mixed sex Canton-S flies exposed to *E. muscae* (961 survivors and 345 zombies). These data were processed into 61-dimensional feature vectors, each representing an individual fly’s behavior up to a particular time of observation. The variables in the feature vector were as follows:

- Feature 1: the time of observation since the start of the experiment (in hours).
- Features 2–11: historical y position values at 10 frames logarithmically spaced between the start of the experiment and 10 minutes prior to the time of observation. frames near the start of the experiment are chosen more sparsely than more recent frames. See Fig 4B.
- Features 12–21: historical fly speed, at the same logarithmically-sampled frames as described above.
- Features 22–41: recent y position at frames uniformly spaced between the time of observation and 10 minutes prior.
- Features 42–61: recent fly speed, at the same uniformly-sampled frames as described above.

Two hundred feature vectors were generated for each fly by selecting 200 random times of observation uniformly distributed across the experiment. Thus, the data set might independently include a feature vector for fly A at ZT13:30 as well as fly at ZT8:00. This yielded a total of 261,200 vectors.

Each feature vector was paired with one of four summiting labels (never-summit, pre-summiting, during summiting or post-summiting). The resultant dataset of 61-dimensional feature vectors and summiting status labels was then randomly subdivided: 75% were used to train a random-forest classifier, and the remaining 25% were withheld as a validation set to evaluate classifier performance. We varied the random forest parameters until satisfactory classifier performance was achieved. At this point, the classifier was tested on a novel experimental dataset generated from a single summiting behavior experiment to assess performance.

In experiments utilizing the classifier in real time, a fly was called as summiting as soon as the predicted during-summiting label probability exceeded the predicted non-infected probability for three consecutive prediction frames (a span of 8 minutes). For experiments requiring paired non-summiting control flies for each flagged summiting fly, five non-summiting candidates were chosen by picking the flies with the highest “non-summiting” score, constructed by multiplying the following four factors:

- the average never-summit label probability over the duration of the experiment
- 1 - the maximum predicted during-summiting probability
- whether the fly was moving at least 10% of all frames in the experiment so far
- the current speed percentile

These factors were chosen heuristically to boost active flies showing few signs of summiting.

### Brain and CA morphology during summiting

Female summiting flies were identified in real time using the random forest classifier, then quickly collected from the summiting assay using a vacuum-connected aspirator and anesthetized with CO2 before being placed on ice. These flies were harvested no earlier than ZT12 on the fourth or fifth day following *E. muscae* exposure. Tissues were dissected and kept ice cold until they were mounted in Vectashield to monitor endogenous fluorescence (in the case of Aug21>GFP flies) or subjected to fixation and subsequent immunohistochemistry (HisRFP and R19G10>mcd8GFP flies).

Corpora allata of Aug21-GFP summiting females were dissected by gently separating the head from the thorax to expose the esophagus and proventriculus. The foregut was severed posterior to the proventriculus and the tissue was mounted in a drop of Vectashield deposited in the middle of three stacked 3-hole reinforcer stickers on a #1 22×22 mm coverslip with the back of the head (posterior side) down. The coverslip was then mounted on an untreated glass slide by gently lowering the slide onto the coverslip until adhesion. The slide was then inverted and imaged at 10x magnification on an upright epifluorescent compound microscope (Nikon 80i) using a constant exposure across samples (300 ms).

Fungal nuclei (HisRFP brains: Hoechst positive, HisRFP negative) or neuropil holes (R19G10>mcd8GFP brains: oval voids) were manually counted in three brain-wide z-stacks (2µm z-step) of HisRFP brains using FIJI (Schindelin et al., 2012). All fungal nuclei were counted in each plane. A comparison of the fraction of nuclei using the manual “raw” method (counting every nucleus across every plane) to an estimate of the actual number of nuclei (via computational collapsing of nuclei counts if their centers are within 2µm in x and y dimensions and 10µm in z) showed both methods gave comparable estimates of the distribution of fungal nuclei across brain regions (Fig 5-S1C, D).Therefore, raw counts were used. Pars intercerebralis cell bodies (R19G10>mcd8GFP brains) were counted in Zen Blue (Zeiss). Each cell body was counted only once, since for this analysis we were investigating the total number of these cells, not their distribution.

### Whole body morphology during end of life

*E. muscae-*exposed Canton-S flies were manually staged at five distinct end-of-life stages and subjected to paraffin embedding, histology, and microscopy in Michael Eisen’s lab at UC Berkeley. Briefly, flies were transferred at 72 hours after exposure to *E. muscae* to individual 500 µL Eppendorf tubes prepared with 100 µL of permissive medium and a ventilation hole poked in the lid with an 18 gauge needle. Flies were manually monitored from ZT8 to ZT13 and immediately immersed in fixative when the following behaviors were first observed: 1) cessation of flight (fly appears to walk normally but does not fly when provoked by the experimenter; corresponds to mid or late summiting), 2) cessation of walking (fly continues to stand upright with proboscis retracted but no longer initiates sustained walking behavior in response to provocation), 3) proboscis extension (proboscis is extended but wings remain horizontal), 4) mid-wing raise (proboscis is extended and wings are approximately half-raised), 5) full-wing raise (proboscis is extended and wings have stopped raising). Paraffin-embedded flies were sliced into eight micron sections and stained with safranin and fast green to visualize interior structures (Elya and Martinez, 2017). Two flies were sectioned for each stage, one sliced sagittally and the other coronally, and imaged on a Zeiss Axio Scan.Z1 Slide Scanner at the Molecular Imaging Center (UC Berkeley).

### Blood-brain barrier integrity

Canton-S flies were exposed to *E. muscae* or housed under mock exposure conditions as previously described. At ZT14 on day four following exposure, ∼50 exposed female flies exhibiting extensive abdominal fungal growth with very white and opaque abdomens (“creamy-bellied”), ∼50 exposed females flies of normal appearance and ∼50 unexposed controls were injected in the mesopleuron with a cocktail of rhodamine B (1.44 mg/mL, Sigma R6626) and 10 KDa dextran conjugated to Cascade Blue (20 mg/mL, Invitrogen D1976) using a pulled glass capillary needle mounted in a brass needle holder (Tritech Research MINJ-4) connected to a 20 mL syringe. The dye cocktail was injected until the anterior abdomen was visibly colored, but not with so much as to completely fill the body cavity and lead to proboscis extension. The volume of injected dye was approximately 75 nL per fly. Injected flies were transferred to foil-wrapped vials containing 5AS to recover. Foil-wrapped vials were placed in an opaque box to further minimize light exposure. After 4 hours, flies were anesthetized with CO2, and their eye fluorescence was scored by an experimenter blind to experimental treatment. Prior to assessing eye fluorescence, flies were screened for rhodamine B fluorescence in the whole body. Flies with weak whole body fluorescence excluded from scoring as they were not loaded with enough dye. Flies were considered “bright-eyed” if there was fluorescence across the entire eye and “dark-eyed” if fluorescence was only apparent at the pseudopupil. Eye fluorescence was used to infer that RhoB was in the brain (Mayer et al., 2009).

### Metabolomics of summiting flies

In two separate experiments, hemolymph was extracted from summiting, non-summiting, and unexposed female flies. In the first experiment, summiting and non-summiting flies were identified manually. This was achieved by releasing *E. muscae*-exposed flies at ∼ZT17 of the third or fourth day following exposure into a large insect rearing cage (Bugdorm #BD4F3030) and continuous visual monitoring of flies from ZT8:30 until ZT11:30 the following day for signs of infection (creamy belly and lack of flight upon provocation). Flies that did not fly and/or right themselves after provoked by the experimenter were designated summiting and collected. For each summiting fly collected, one exposed fly that did respond to provocation (non-summiting) and one unexposed fly (kept in a separate enclosure, unexposed) were collected simultaneously. All flies were retrieved from their enclosures using mouth aspiration, then stored on ice in Eppendorf tubes until a total of 20 flies had been collected. This was repeated to obtain duplicate pools of 20 flies for each infection status (summiting, non-summiting and unexposed).

For the second experiment, summiting and non-summiting flies were identified in real-time using the random forest classifier. *E. muscae*-exposed females were loaded into standard summit arenas on the third or fourth evening following *E. muscae* exposure and tracked until ZT13 of the following day. Summiting and non-summiting flies were flagged in pairs automatically and the experimenter was alerted by email. Flies were promptly collected using a vacuum-assisted aspirator then briefly anesthetized with CO2 and placed in 1.7 mL Eppendorf tubes on ice until twenty individuals were collected per treatment. An unexposed control fly was collected simultaneously with every summiting/ non-summiting pair. Triplicate pools of 20 flies were collected for each infection status.

Hemolymph was extracted from a pool of 20 flies by piercing the mesopleuron of each with a 0.2 Minutien pin (Fine Science Tools) mounted on a nickel plated pin holder (Fine Science Tools) under CO2 anesthesia (Musselman, 2013). Pierced flies were transferred to a 500 µL microcentrifuge tube pierced at the bottom with a 29 1/2 gauge needle nested in a 1.7 mL Eppendorf tube. Tubes were centrifuged at room temperature for 10 min at 2700g to collect a droplet of hemolymph. Hemolymph was stored on ice until all samples had been extracted. Samples for metabolomic analysis were 1 µL of hemolymph added to 2 µL of 1x PBS.

Metabolite detection and putative compound identification were performed by the Harvard Center for Mass Spectrometry. Hemolymph samples were brought to a final volume of 20 µL with the addition of acetonitrile, to precipitate proteins. Following centrifugation, 5 µL of supernatant was separated on a SeqQuant Zic-pHILIC 5 um column (Millipore #150460). For each experiment, solvent mixtures comprising 20 mM ammonium carbonate, 0.1% ammonium hydroxide in water (solvent A) and 97% acetonitrile in water (solvent B) were flowed for ∼50 minutes at 40°C. For the manually-staged experiment, the following solvent mixtures were flowed at 0.2 mL per minute: 100% B (20 min), 40% B, 60% A (10 min), 100% A (5 min), 100% A (5 min), 100% B (10 min). For the classifier-staged experiment, the following solvent mixtures were flowed at 0.15 mL per min: 99% B (17 min), 40% B + 60 % A (10 min), 100% A (5 min), 100% A (4 min), 99% B (11 min). For the manually-staged experiment, separated compounds were fragmented using electrospray ionization (ESI+) and detected using a ThermoFisher Q-exactive mass spectrometer under each positive and negative polarity (Resolution: 70,000, AGC target: 3e6, mz range: 66.7 to 1000). For the classifier-staged experiment, separated compounds were fragmented using heated electrospray ionization (HESI+) and detected using a ThermoFisher Orbitrap ID-X mass spectrometer under each positive and negative polarity (Resolution: 500,000, AGC target: 1e5, mz range: 65 to 1000). The variations in flow rate and ionization protocol were unlikely to substantially affect the compounds we were able to detect between the experiments.

MS-MS was performed twice (once each for the manual and classifier-staged experiments) on mixed pools (5 µL of each of the three samples per experiment)) using AcquireX DeepScan in each positive and negative modes and 2 level depth. All data were normalized (median centering) to compensate for biomass differences and analyzed with Compound Discoverer v. 3.1 (ThermoFisher Scientific). Molecular formulae were predicted from measured mass and isotopic pattern fit. Abundance values were determined for every peak observed within the MS-MS experimental pool for every sample. All chromatograms were manually checked to distinguish likely real signal from noise, with compounds typically considered absent from a sample if intensity counts were <1e3. Putative compound identities were manually assigned from high-confidence database matches (MZcloud, MZvault, HCMS locally-curated mass list) based on accurate mass and MS-MS spectra. Compounds were considered to be observed in both experiments (manually-staged and classifier-staged) if their molecular weights were within 5 ppm. All MS data are available in File S1.

### Hemolymph transfusion

Three and four days prior to the transfusion experiment, mixed sex Canton-S flies were exposed to *E. muscae* in cages as described above. One day prior to the transfusion experiment, flies destined to receive hemolymph (either unexposed Canton-S males or 72 hour exposed Canton-S females; Fig 6C, D) were transferred into individual housing consisting of PCR tubes containing ∼100 µL of 5AS and with two holes poked in the cap using an 18 gauge needle to provide airflow. Donor flies (females exposed ∼72 or ∼96 hours prior) were loaded into standard summiting arenas. Donor tracking began at ∼ZT17 and the summiting classifier was launched.

The next day, two experimenters, A and B for the purposes of this explanation, implemented the transfusion experiment from ZT8 until ZT12 or until 32 pairs of recipient flies had been transfused. Experimenter A collected donor flies; experimenter B performed the transfusions. Each transfusion began when a fly was flagged by the classifier (see *Real-time summiting classifier)* and Experimenter A was alerted via email. Experimenter A inspected behavioral traces to confirm the accuracy of the summiting classification and selected one of five identified non-summiters to serve as a time-matched control. Experimenter A then collected these flies from arenas via vacuum-assisted aspiration, anesthetized them with CO2, and placed them in adjacent wells of a 96 well plate on ice. Fly placement was randomized (i.e., sometimes the summiting fly was placed first, sometimes the non-summiting fly) and recorded before the plate was passed to Experimenter B. Thus, Experimenter B was blind to fly summiting status. Experimenter B then used a pulled capillary needle to remove ∼50 nL of hemolymph from the first donor through the mesopleuron and injected this material into the mesopleuron of a cold-anesthetized recipient. The needle was rinsed thoroughly in molecular grade water between transfusions. Immediately after transfusion, recipient flies were transferred to standard summiting arenas that were already in place in an imaging box and being tracked by MARGO. Tracking continued for no less than three hours after the final fly had been transfused. Used donor flies were transferred into individual housing and monitored for the next 48 hours for death by *E. muscae*.

Behavioral data were processed blind by Experimenter B. The time of recovery from anesthesia (i.e., resumption of locomotion) was manually determined for each fly based on its behavioral trace. Flies that did not recover or showed very little total movement were discarded from subsequent analysis. Fly summiting status was then revealed by Experimenter A to determine average distance traveled vs time for each treatment group. After the data had been curated in this blinded manner, Experimenter A revealed the behavior calls for each donor to Experimenter B. Experimenter B used this information as well as donor outcome to determine the average distance traveled for each treatment group. Donors that were identified as summiting but failed to sporulate on the day of the experiment were interpreted as misclassified and their corresponding recipients were dropped from the analysis.

## Acknowledgements

We thank Ryan Maloney and David Zimmerman for helpful comments on the manuscript. We are indebted to Ed Soucy and Brett Graham of Harvard University’s Center for Brain Science’s magnificent Neuroengineering Core for their help in designing and fabricating the summiting behavior assay and to Charles Vidoudez of the Harvard Center for Mass Spectrometry for generating and curating hemolymph metabolomic data. We are also grateful to many folks for generously sharing reagents: Rochele Yamamoto (*Aug21-Gal4, UAS-GFP* flies), David Anderson (*UAS-mCherry*.*FRT*.*eGFP-Kir2*.*1* a n d *UAS-eGFP-Kir2*.*1*.*FRT*.*mCherry* flies), Daniel Cavanaugh (*VTDh44-Gal4, Clk856-Gal4*, and pan-DN1p-*Gal4* flies), Matthias Schlichting (*UAS-Cas9, UAS-PdfR-sgRNA* flies), Toshihiro Kitamoto (*Cha-Gal80* flies), Jess Kanwal (*UAS-Kir2*.*1* flies), Kristin Scott (*nan<sub>36a </sub>*and *tub-Gal80<sub>ts </sub>*flies), Kendal Broadie (*tutl<sub>1 </sub>*flies), and Ryusuke Niwa (JHAMT primary antibody). We also would like to thank all of the core facilities that enabled this work: the Harvard Center for Mass Spectrometry (LC-MS), the Harvard Center for Biological Imaging (confocal imaging), UC Berkeley’s Biological Imaging Facility (whole fly microtomy and histology), UC Berkeley’s Molecular Imaging Center (whole fly histology imaging), the Kyoto Drosophila Stock Center (fly lines), Janelia Research Campus (fly lines), Bloomington Drosophila Stock Center (many, many fly lines). Finally, we are grateful to all of the hard-working individuals (i.e., *E. muscae*teers) who have helped keep the fungus thriving in the lab: Benno Rodemann, Fosca Bechthold, Aundrea Kroger, Nicole Pittoors, Ryan Maloney, Stanislav Lazopulo, Haesung Jee, Dylan Roy and Noah Rodman.

## Funding

CE was supported by a Harvard Mind, Brain and Behavior post-doctoral fellowship and a HHMI Hanna H. Gray postdoctoral fellowship; DL was supported by the NSF-Simons Center for Mathematical and Statistical Analysis of Biology at Harvard, award number #1764269 and the Harvard Quantitative Biology Initiative; BdB was supported by a Sloan Research Fellowship, a Klingenstein-Simons Fellowship Award, a Smith Family Odyssey Award, a Harvard/MIT Basic Neuroscience Grant, National Science Foundation grant no. IOS-1557913, and NIH/ NINDS grant no. 1R01NS121874-01.

## Conflicts of Interest

The authors declare no competing interests.

## Supplementary materials

**Figure 1-S1.**
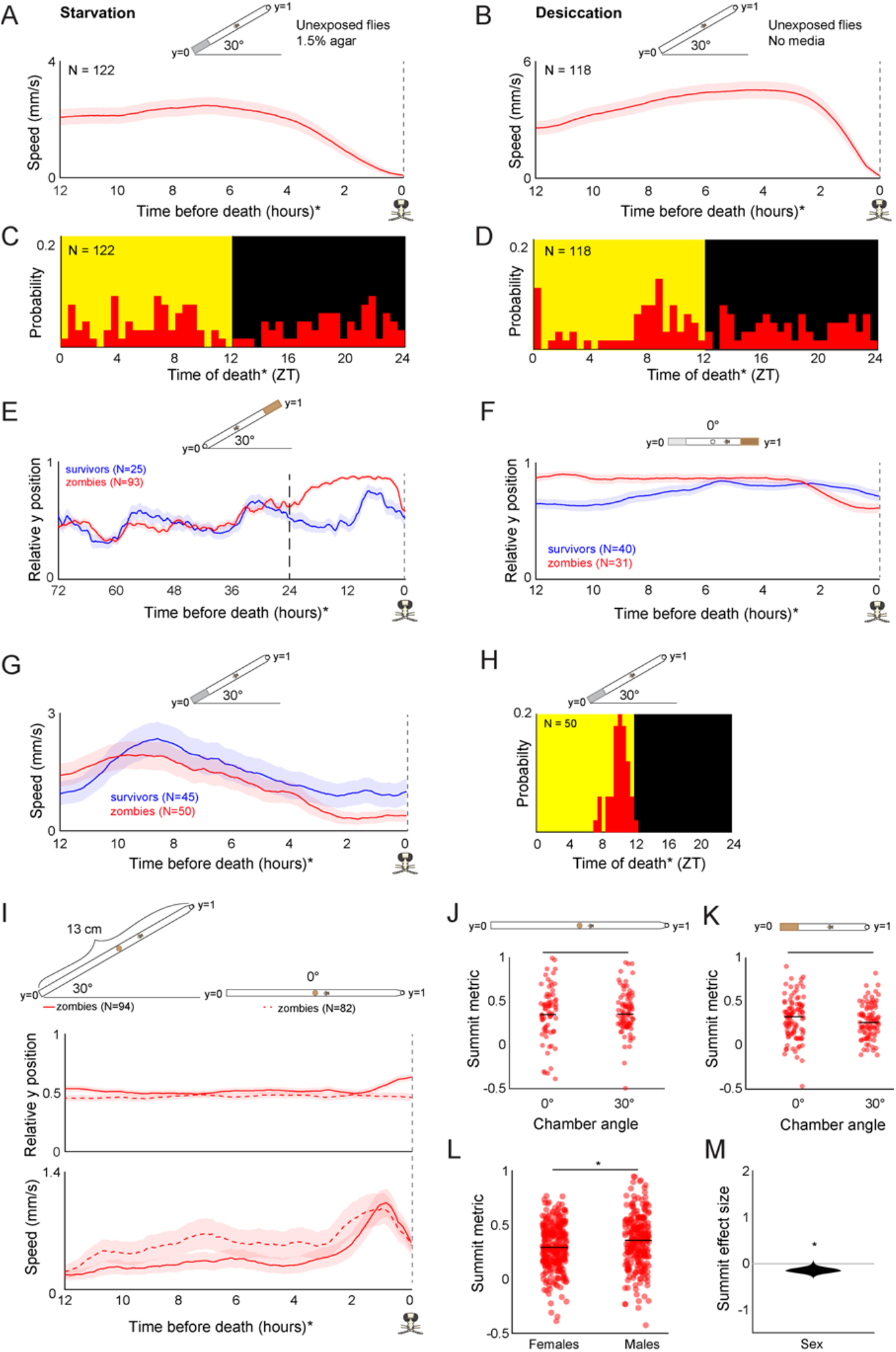
Additional features of summiting behavior in the custom behavior assay. A) Unexposed, wild-type flies that die of starvation do not exhibit a burst in activity beginning 2.5 hours prior to their last movement and C) do not exhibit specific circadian timing of death. Flies in this experiment were provided non-nutritive agar instead of food at position y = 0. B) Unexposed, wild-type flies that die of desiccation do not exhibit a burst in locomotor activity beginning 2.5 hours prior to their last movement and D) do not exhibit specific circadian timing of death. Flies in this experiment were provided food nor agar at position y = 0. E) Mean y position of zombie flies diverges from survivors beginning around 24 hours prior to death when nutritive food is positioned at y = 1. F) Compared to survivors, zombie flies prefer nutritive (5% sucrose, 1.5% agar) over non-nutritive (0% sucrose, 1.5% agar) media in their final 12 hours of life prior to the onset of summiting at 2.5 hours before death. G) Zombie flies housed on non-nutritive food for their final 24 hours do not exhibit summiting behavior and H) show typical timing of last death (compare to Fig 1E). I) When housed in tall arenas (13 cm) with food placed in the middle, zombie flies tend to move slightly upward when the arena is tilted at 30°, but not 0°. Zombies in arenas at both angles exhibit a burst of speed prior to death (bottom). J, K) Summit metric (see Fig 1I) does not vary for zombies housed in chambers at 0° versus 30° in tall or standard chambers. L) Summit metric for female versus male Canton-S zombies. M) Distribution of estimated effect size of sex on summiting. For A, B, E-K, diagrams depicting behavior arena setup are shown above corresponding plots. For L and M, behavior arenas were in standard configuration (as in Fig 1F)* = p<0.05; ** = p<0.01; *** p<0.001 by two-tailed t-test.

**Figure 2-S1.**
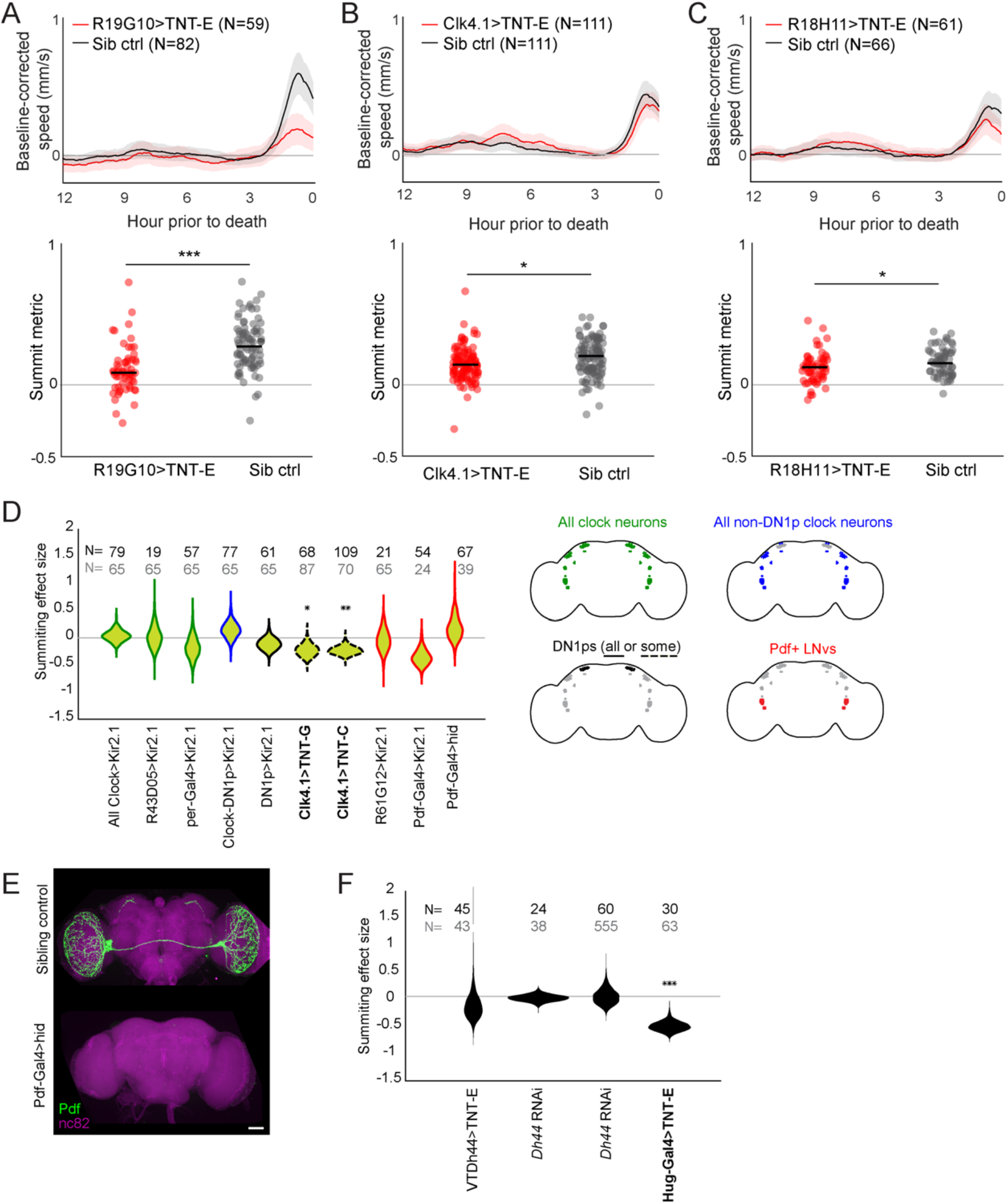
Additional experiments assessing summiting after clock neuron and R19G10 disruption. A-C) Plots visualizing summiting for manipulations targeting R19G10 and DN1p neurons. Top: Mean zombie baseline-corrected speed versus time. Shaded area is + / - one standard error. Bottom: Individual SM (circles) and median SM (line). * = p<0.05; ** = p<0.01; *** p<0.001 by two-tailed t-test. Clk4.1 drives expression in 8-10 and R18H11 drives expression in 7-8 of DN1p neurons per hemisphere, respectively (Kunst et al., 2014; Zhang et al., 2010). D) Left: Summiting effect size estimate distributions for Gal4/effector combinations targeting clock neurons. Right: Diagrams depicting target cells of tested reagents following Shafer et al., 2022. Violin plot outline colors match diagrams. * = p<0.05; ** = p<0.01; *** p<0.001 by two-tailed t-test. E) Confocal micrographs demonstrating ablation of Pdf-expressing neurons in Pdf-Gal4>hid animals. F) Summiting effect size estimate distributions when targeting components of the circadian locomotor output pathway identified by (Cavanaugh et al., 2014). * = p<0.05; ** = p<0.01; *** p<0.001 by two-tailed t-test.

**Figure 2-S2.**
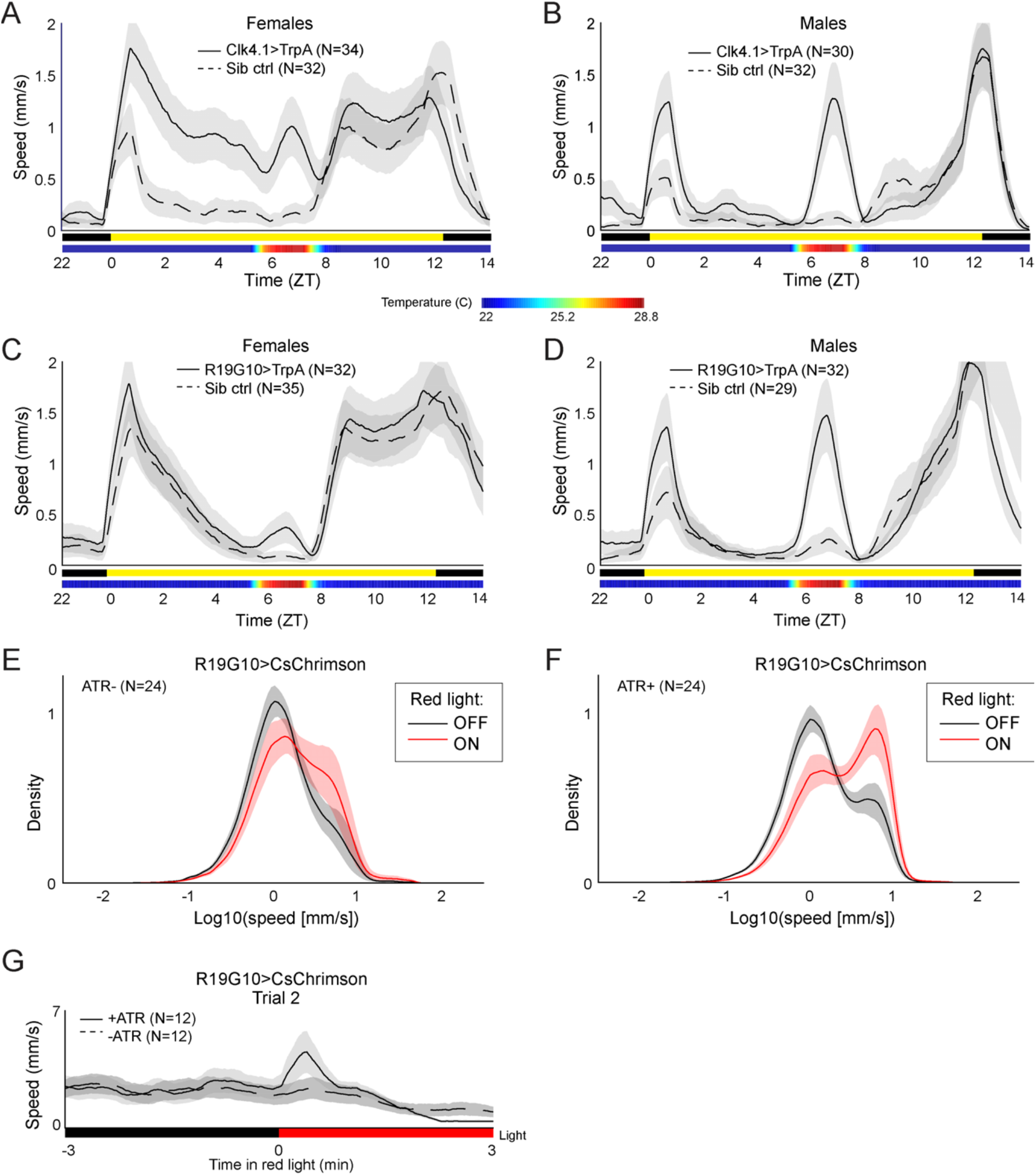
Additional experiments assessing sufficiency of DN1p and R19G10 neuron activation for increased locomotion. A-D) Mean speed of unexposed flies with thermogenetically activated DN1p or R19G10 neurons (solid line) versus sibling controls (dashed line) separated by sex. The presence or absence of light is indicated below the horizontal axis with a yellow or black bar, respectively. The measured temperature of the room is shown as a heat map below the light cue indicator. Shaded regions are +/- one standard error. For A and B, Clk4.1-Gal4 was used to drive expression in DN1ps. E & F) Kernel density estimates of the distribution of log-speed for R19G10 > CsChrimson flies with (F) or without (E) of dietary ATR in the presence of absence of 5 Hz pulsed red light stimulus (OFF and ON, respectively). A bimodal distribution of speeds is observed regardless of ATR and light treatment. The lower peak (centered around 0) corresponds to non-walking and the higher peak (centered around 6mm/s) corresponds to walking flies. G) Mean speed across flies of unexposed R19G10>CsChrimson flies before and after the second presentation of red light in the experiment described for Fig 2K. ATR+ flies show smaller increase in locomotion upon red light stimulus, consistent with CsChrimson depolarization block or adaptation within the stimulated circuit.

**Figure 3-S1.**
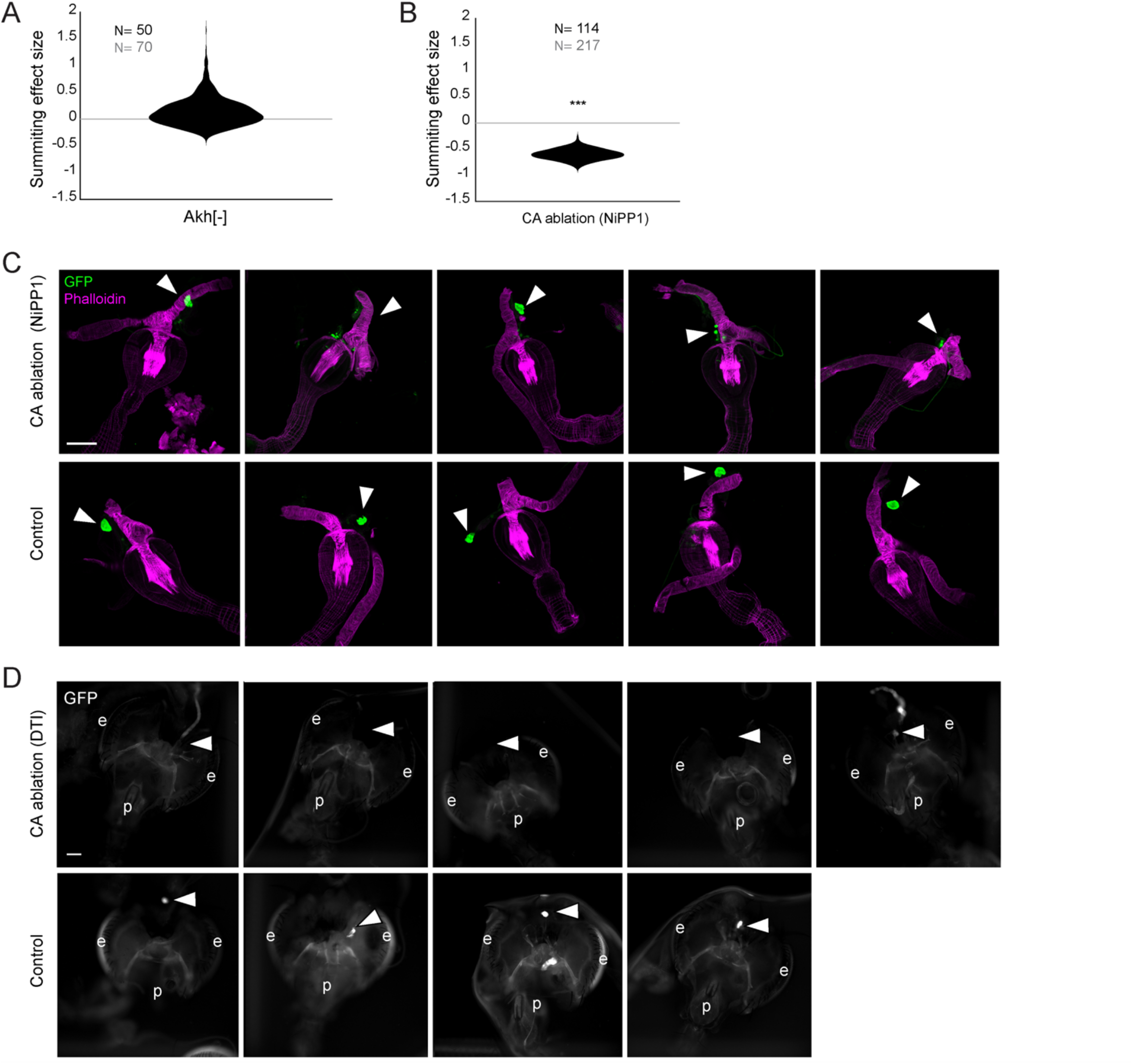
Supporting data for juvenile hormone involvement in summiting. A) Summiting effect size estimate distribution for *Akh-*mutants, showing no effect compared to controls. Sample size of experimental animals in black, controls in gray. * = p<0.05; ** = p<0.01; *** p<0.001 by two-tailed t-test. B) As in (A) for the ablation of the CA using *NiPP1* effector (per Yamamoto et al, 2013). A significant reduction in summiting is seen compared to controls. C) Confocal micrographs of the anterior foregut and retrocerebral complexes from Aug21>GFP, NiPP1 flies and Aug21>GFP sibling controls. Green channel is GFP and magenta anti-actin phalloidin counterstain. Arrowheads indicate observed or expected location of CA. Scale bar is 100 microns. CAs in Aug21>GFP, NiPP1 animals ranged from intact to absent. D) Compound epifluorescence micrographs of the head, anterior foregut and retrocerebral complexes of tub-Gal80, Aug21> GFP, DTI flies. In animals in which DTI expression was induced by thermal inactivation of the Gal80 repressor, only one sample showed potential residual CA; all other animals lacked CA.CA were observed in all controls. e = eye, p = proboscis.

**Figure 3-S2.**
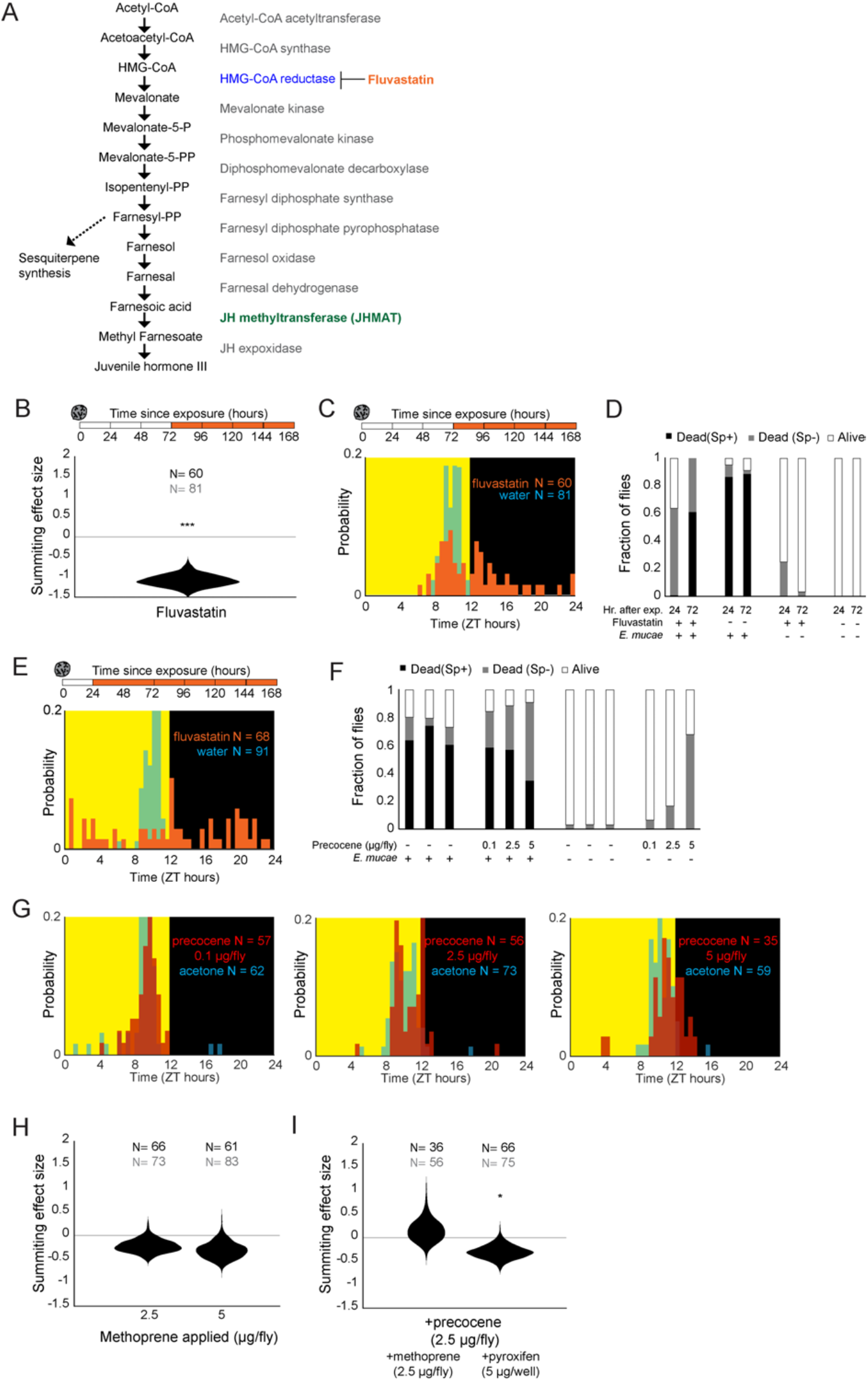
Additional experiments examining juvenile hormone involvement in summiting. A) Synthesis pathway for juvenile hormone. Enzymes catalyzing each step are shown at right. HMG-CoA reductase (blue) is the target of the drug fluvastatin. B) Summiting effect size estimate distribution for *E. muscae*-exposed flies fed with fluvastatin (250 µg/well) starting 72 hours after exposure to *E. muscae*. * = p<0.05; ** = p<0.01; *** p<0.001 by two-tailed t-test. Sample size of experimental animals in black, controls in gray. C) Times of death for flies fed fluvastatin 72 hours after exposure and killed by *E. muscae* (i.e., with sporulation). D) Stacked bar plots of survival outcomes for fluvastatin-treated flies and controls, at 72 hours and 24 hours after *E. muscae* exposure. Flies were manually assessed as alive or dead at the end of behavior tracking; dead flies were further classified as having sporulated (Sp+) or not (Sp-). *E. muscae*-exposed N = 91-108; unexposed N = 20-37. E) Times of death for flies fed fluvastatin 24 hours after exposure, which showed no signs of sporulation and died roughly uniformly throughout the day. F) Stacked bar plots of survival outcomes for flies treated topically with indicated amount of precocene or vehicle control (acetone) with or without exposure to *E. muscae. E. muscae*-exposed N = 97-100; unexposed N = 28-31.G) Times of death for flies exposed to *E. muscae* and treated with three concentrations of precocene or acetone at 72 hours after exposure. H) Summiting effect size estimate distributions for topical application of the JH analog methoprene at two different doses. This had no effect compared to vehicle (acetone) controls. I) Summiting effect size estimate distributions for co-applied methoprene or dietary pyriproxyfen. These treatments did not rescue summiting deficits induced by precocene.

**Figure 4-S1.**
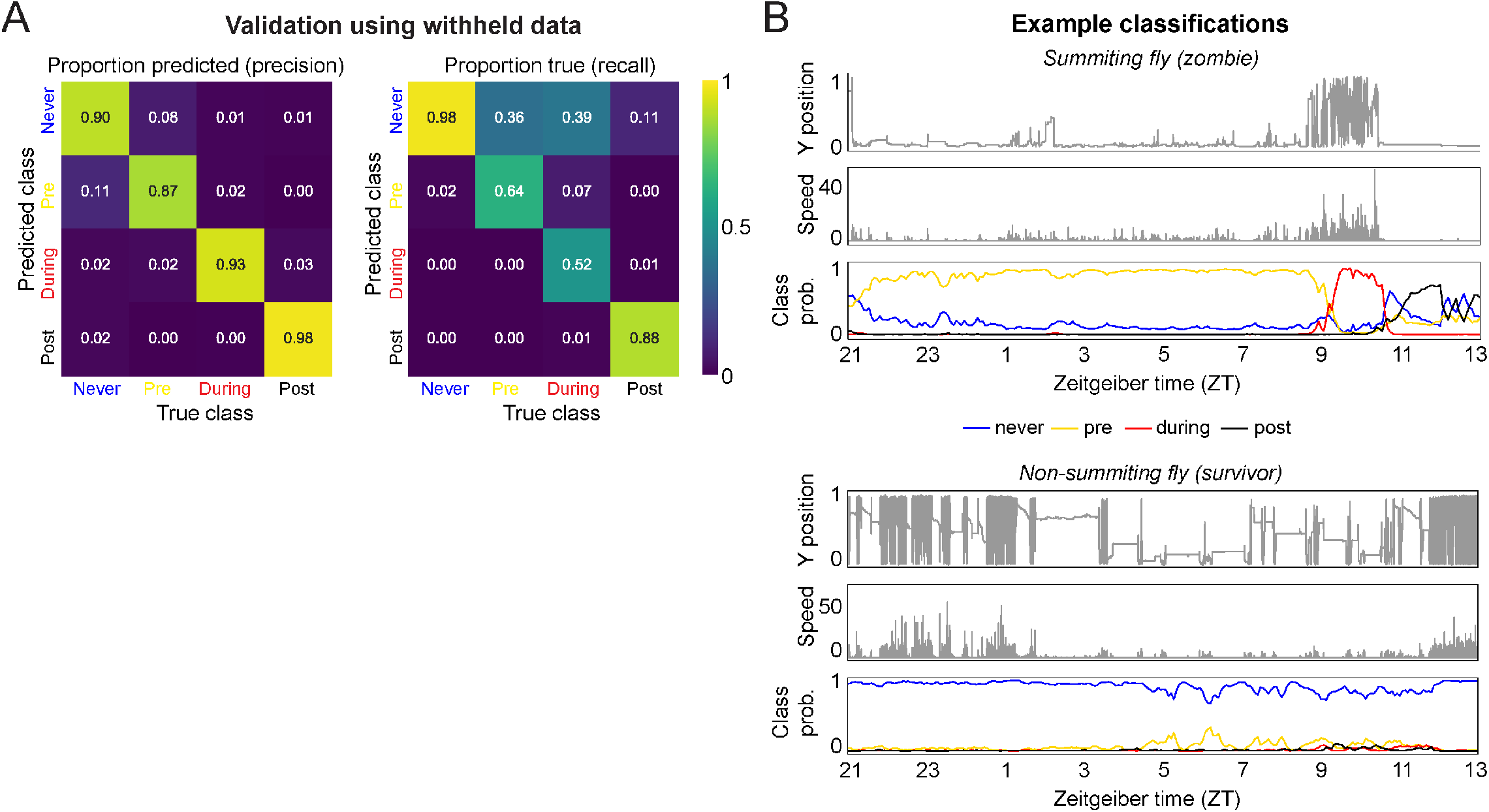
Development of a real-time random forest classifier for summiting behavior. A) Confusion matrices calculated using the validation data set (25% of the total ground truth data). B) Example classifications of a summiting (zombie) and a non-summiting (survivor) fly over an entire behavior tracking experiment. Top and middle plots for each example are the fly’s y position and speed; bottom plot is the class probabilities for the fly at each timepoint. In the summiting example, the fly is consistently classified as pre-summiting (i.e., will become a zombie before the next occurrence of sunset) starting as early as ZT22 the evening prior to death. This fly was classified as summiting from approximately ZT9.25 to ZT10.75 the following day, and post-summiting (i.e., dead) ZT11 onward. In contrast, the non-summiting example fly was consistently classified as never-summiting (i.e., would live through the next sunset) for the duration of the experiment.

**Figure 5-S1.**
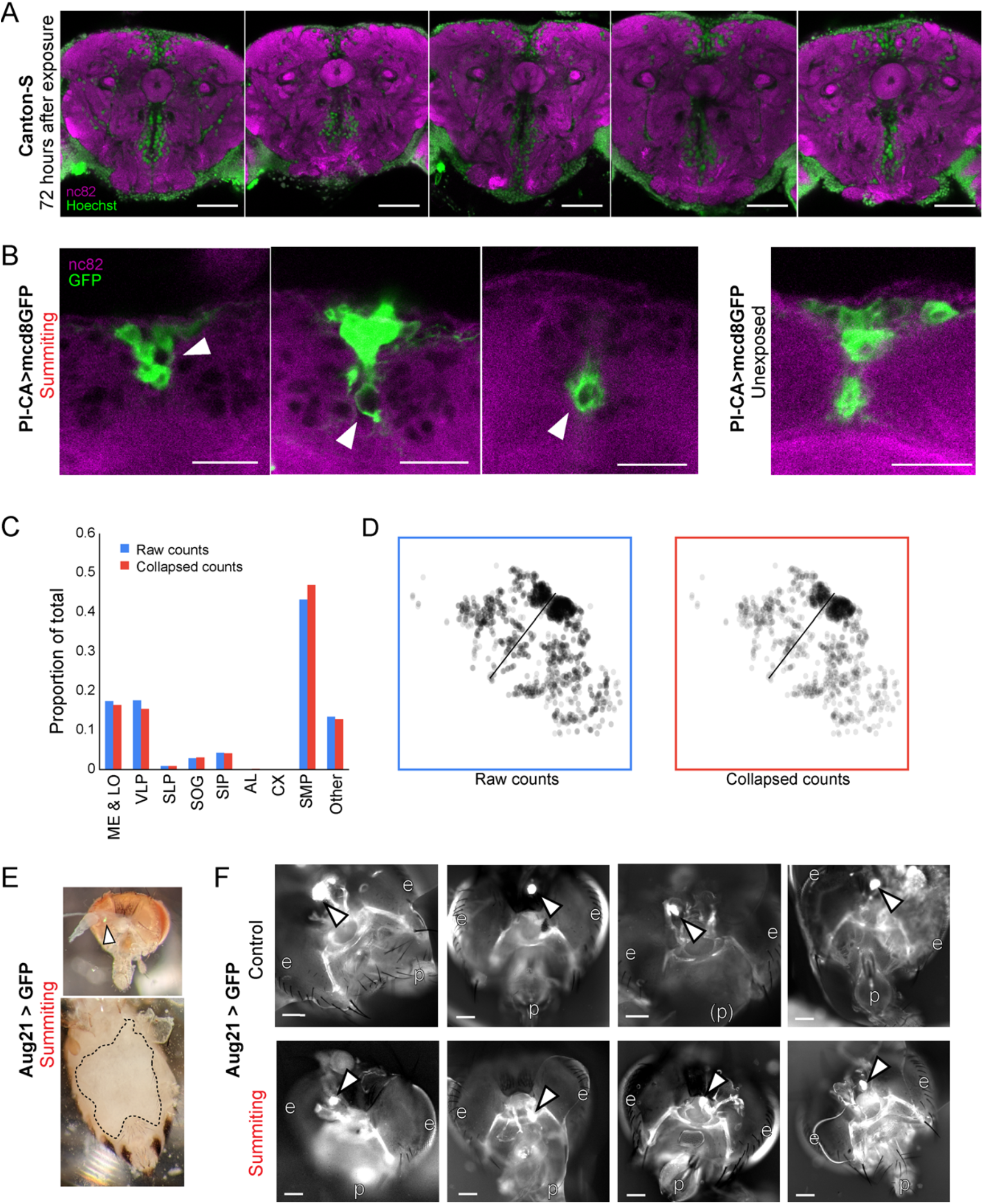
Supporting data for host morphology during *E. muscae* infection. A) Confocal micrographs of five infected brains 72 hours after *E. muscae* exposure. Green is Hoechst and marks all nuclei. Magenta is the synaptic marker nc82. Fungal cells are present in the SMP and absent from the CX. Scale bar is 50 microns. B) Confocal micrographs if PI-CA>mcd8GFP (green) brains during summiting and unexposed to *E muscae*. White arrowheads indicate apparent displacement of PI-CA processes by fungal cells. Scale bar = 25 microns. C) Distribution of *E. muscae* nuclei across brain regions from a whole-brain confocal volume (2 micron z-step), using two counting methods. Counting every nucleus seen in a single z-slice as one cell (“Raw counts”), and merging cell bodies if they have the same x- and y-position (to within 2 µm) and appear within 10 microns of each other along the z-axis (“Collapsed counts”) produces very similar distribution estimates (N = 1,426 nuclei). D) Z-projections of nuclei as counted by the Raw and Collapsed methods. E) Micrographs of dissected summiting Aug21>GFP fly. Top: Head (brightfield overlaid with GFP channel). White arrow points to the CA. Bottom: Abdomen (brightfield) with ventral cuticle peeled back. Dashed line indicates the edges of the fly’s remaining cuticle. Only fungal tissues, and no host organs are visible inside the dashed line. F) Additional whole-mount preparations of Aug21>GFP heads and retrocerebral complexes in control (non-summiting) and summiting flies, as in Fig 5. White arrows indicate CA; p=proboscis, e=eyes. Scale bars each 100 microns.

**Figure 6-S1.**
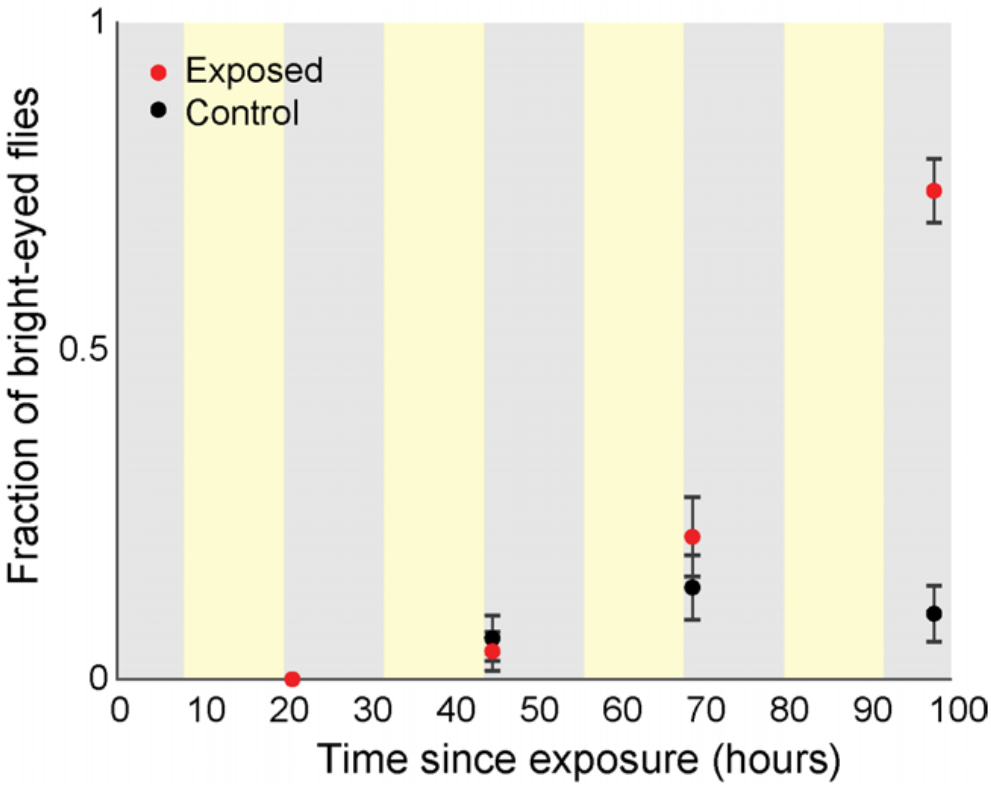
blood-brain permeability as a function of time since exposure. Flies were injected with 1.44 mg/mL RhoB then allowed to incubate for four hours prior to scoring eye fluorescence. Experimenters were blind to treatment type. Error bars are the estimated standard error of the binomial proportion. Background shading indicates entrained lighting conditions experienced by all flies (yellow = lights on, gray = lights off).

**Figure 6-S2.**
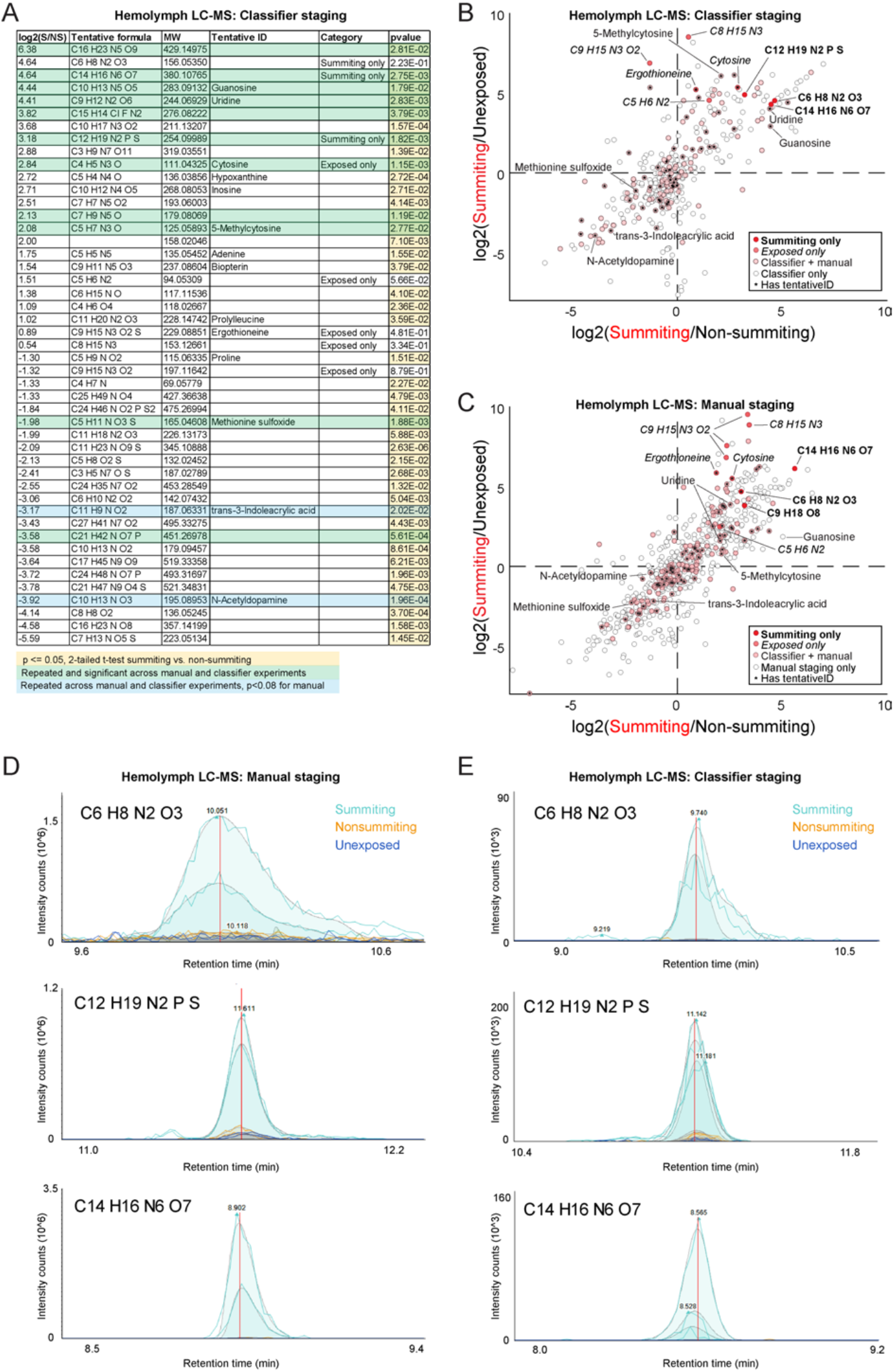
Metabolomics of summiting flies. A) Compounds of interest from classifier-staged metabolomics experiment. Compounds are included if at least one of three conditions is met: 1) they are significantly different between summiting and non-summiting, exposed animals, and exhibit at least a 2-fold change between these treatments, 2) they were only detected in summiting flies (“Summiting only”), or 3) they were only detected in exposed flies (“Exposed only”; summiting or non-summiting), but not unexposed flies. P-values <0.05 are highlighted in yellow. Compounds highlighted in green were observed to be significantly different between summiting and non-summiting flies in both the manual and classifier-based experiments (p <= 0.05 by a two-tailed t-test); compounds in blue were significant in the classifier-staged experiment and fell just short of significant (p<0.08) in the manual experiment. B) Scatter plot of the log2-fold-change of compound abundance in summiting compared to unexposed flies versus the log2-fold-change of compound abundance in summiting compared to exposed, non-summiting flies. Data from the classifier-staged experiment. Compounds in the darkest red were observed only to have chromatogram peaks above baseline (1e3 intensity counts) in summiting samples and not in any other samples. Compounds in medium red were only observed to have real peaks in exposed (summiting or non-summiting) but not unexposed samples across both experiments. Compounds in light red were observed in all experimental groups of both experiments (molecular weight within five parts per million between experiments). Compounds in white were observed in the classifier-based experiment but were not observed in the manually-staged experiment (absolute difference in molecular weight greater than five ppm). Compounds that were observed in both experiments, showed significant differences between summiting and non-summiting flies, and were tentatively identified, are labeled with compound name or molecular formula. C) As in (B), for the manually staged experiment. D,E) Raw chromatogram peaks for three compounds assessed to only be present in summiting flies in manually-staged (D) and classifier-staged (E) metabolomics experiments. Different peaks of the same color are biological replicates. Putative formulas as determined from the measured mass given in the upper left corner for each plot.

**File S1. Metabolomics data for both manual and classifier-assisted experiments**. *http://lab.debivort.org/zombie-summiting/Elya_et_al_2022_summiting_MSMS_data.xlsx* — Sheets with prefix “All_” contain all of the data from the indicated metabolomics experiment. Sheet with prefix “Overlap_” contains data for compounds that were observed in both experiments (compounds whose molecular weight corresponded within 5 parts per million [ppm]). Column headers ending in (M) indicate results from the manually-staged experiment; (C) from the classifier-staged experiments. Annotations for columns A, B, C, K, and N are as follows (with values of 1 or TRUE indicating the condition is met, 0 or FALSE that the condition is not met): “Significant” = significantly different between NS and S across both experiments; “Infected-specific” = compound only present in NS and S, but not C, samples, per manual chromatogram inspection; “Summiting-specific” = compound only present in S, but not NS or C, samples, per manual chromatogram inspections; “Same tentative formula?” = predicted formulae for given compound match exactly across experiments; “Same sign?” = log2 fold change of compound abundance between S and NS samples is consistently positive or negative across experiments. Abbreviations: S = summiting, NS = exposed but non-summiting, C = unexposed control.

## Notes

http://lab.debivort.org/zombie-summiting/

### Competing Interest Statement

The authors have declared no competing interest.

